# Three cortical streams for somatosensory information processing

**DOI:** 10.1101/2025.04.14.647517

**Authors:** Meiqi Niu, Seán Froudist-Walsh, Yujie Hou, Lucija Rapan, Nathan Vinçon, Henry Kennedy, Ting Xu, Nicola Palomero-Gallagher

**Author notes:** Corresponding author: Nicola Palomero-Gallagher Institute of Neuroscience and Medicine (INM-1) Research Centre Jülich 52425 Jülich Germany. Corresponding author: Meiqi Niu Institute of Neuroscience and Medicine (INM-1) Research Centre Jülich 52425 Jülich Germany.

## Abstract

The somatosensory cortex processes information hierarchically, transforming sensory input into appropriate responses. This hierarchy, in turn, provides a fundamental principle for the organization of anatomical and functional properties across the somatosensory cortex. While the local somatosensory hierarchy has been studied, a comprehensive model that fully illustrates somatosensory information transmission in fine detail remains lacking. In this study, we examine multimodal connectivity patterns of the entire macaque somatosensory cortex by integrating the information from receptor covariance (RC) and structural (SC) or functional connectivity (FC). Our findings not only reveal the hierarchical relationships but also propose a model of somatosensory processing streams. In this model, area 3bl serves as the initial cortical stage for somatosensory signals, projecting to areas 3al, 1, and 2. From there, somatosensory signals follow three major pathways: ventrally to the SII complex, medially to the medial SI and TSA, and posteriorly to somatosensory association areas in the parietal lobe. Further analysis shows that RC is not only closely linked to SC and FC but in addition displays unique characteristics that likely relate to the hierarchical processing across sensory modalities. This study deepens our understanding of brain connectivity patterns across different modalities and links the structural, chemoarchitectonic, and functional organization of the macaque somatosensory cortex.

## Introduction

The somatosensory cortex plays a fundamental role in processing sensory information and generating appropriate responses. Its hierarchy has been used to exemplify the systematic organization of the brain (Delhaye et al., 2018; Felleman and Van Essen, 1991; Rossi-Pool et al., 2021). Cortical somatosensory processing hierarchy starts with the primary somatosensory cortex (SI), which processes basic tactile information, followed by the secondary somatosensory cortex (SII) which integrates more complex sensory data. At the highest level, the posterior parietal cortex is responsible for higher-level processing and spatial awareness (Iwamura, 1998; Rossi-Pool et al., 2021). Each one of these “major” hierarchical levels is composed of areas that exhibit diversity in their anatomical features, coding dynamics and functional roles (Kaas, 2012; Niu et al., 2024; Qi et al., 2008; Saadon-Grosman et al., 2020; Thomas et al., 2021). Interestingly, there is no consensus concerning the hierarchical organization within each of these “major” levels, and to our knowledge, no comprehensive model has yet been developed to fully illustrate the flow of somatosensory information transmission.

These areas are intricately woven into a complex network of anatomically connected and functionally interacting neuronal populations and engage in dense interactions to produce and modulate dynamic brain functions (Thiebaut de Schotten and Forkel, 2022). The structural connectivity (SC) of somatosensory areas in monkeys has been extensively studied using retrograde tracer techniques. In SI, area 3b receives somatosensory input from the thalamus and projects to areas 1, 2 and S2 (Burton and Fabri, 1995; Darian-Smith et al., 1993), while area 3a also receives thalamic input but sends projections to area 2, the motor cortex, S2, and the insula (Huffman and Krubitzer, 2001), areas 1 and 2 integrate inputs from 3b, 3a, and thalamus, sending outputs to multiple regions, such as areas 2, S2, PV, the insula, and the posterior parietal cortex (Cusick et al., 1985; Darian-Smith et al., 1993). Areas within SII mainly receive projections from SI areas and thalamus, and project to the areas located in the anterior part of the posterior parietal lobe (Disbrow et al., 2003). As information moves up the somatosensory hierarchy, it becomes progressively processed and integrated. While traditional tracer studies have provided valuable insights into anatomical connectivity, they are technically limited in providing a comprehensive and systematic understanding of connectivity patterns across the entire somatosensory cortex.

Functional MRI (fMRI), especially resting-state fMRI, presents a non-invasive *in-vivo* approach for studying connectivity across different brain regions (Bijsterbosch et al., 2017). Analysis of time series correlations between brain areas can unveil functional connectivity (FC), even in the absence of explicit tasks (Biswal et al., 1995; Van Den Heuvel and Pol, 2010). In the macaque somatosensory cortex, (Thomas et al., 2021) revealed that different body part representations in area 3b have distinct FC patterns, and (Wang et al., 2013) reported strong same-digit interactions between areas 3b and 1, as well as prominent interdigit connections within area 3b. Although previous studies (Fox et al., 2005; Wang et al., 2013) suggested substantial overlap between functional and anatomical connectivity patterns, the neuronal basis underlying the structure-function relationship remains unclear. Specifically, mechanisms governing how resting-state FC is constrained by anatomical connectivity and the interaction between structure and function at the system level, remains to be clarified.

Neurotransmitters and their receptors are the key molecules that enable signal transmission and neuronal communication (Palomero-Gallagher et al., 2015). Receptors of the classical transmitter systems are heterogeneously distributed across the different brain areas, and the well-tuned balance between different receptor types within a given brain area plays a pivotal role in enabling and modulating brain functions (Palomero-Gallagher and Zilles, 2018; Zilles and Palomero-Gallagher, 2017b). In recent years, significant efforts have been dedicated to creating a comprehensive, receptor-driven multimodal atlas of the macaque brain (MEBRAINS atlas; Impieri et al., 2019; Niu et al., 2020; Niu et al., 2024; Niu et al., 2021; Palomero-Gallagher et al., 2013; Pérez-Santos et al., 2021; Rapan et al., 2021; Rapan et al., 2023; Rapan et al., 2022), which provides an opportunity to explore receptor covariance patterns (RC; i.e., receptor similarity pattern) across brain regions. By quantifying RC, we can infer the likelihood of each pair of areas being similarly influenced by endogenous or exogenous input (Hansen et al., 2022), thereby providing insights into the chemoarchitectonic mechanisms of the underlying cortical network. Since the MEBRAINS atlas not only provides information on the distribution of 14 receptors for classical neurotransmitters but also includes data on cell bodies and myelin fibers, its integration of all this information into stereotaxic space enables comparisons with functional datasets. Therefore, using this atlas to explore the RC across areas of interest holds promise as a viable approach for linking brain structural and functional connectivity patterns at both system-wide and fine-grained local cortical scales.

In the present study, we aim to investigate the multimodal covariance of the macaque somatosensory cortex by integrating *receptor covariance*, *structural connectivity*, and *functional connectivity* (Fig. 1). Specifically, we seek to characterize RC patterns of the somatosensory cortex in the macaque monkey brain to examine similarities in chemoarchitectonic balance across brain regions. We hypothesize that variations in multimodal covariance across somatosensory areas will reflect the hierarchical organization or information processing pathways of the macaque somatosensory cortex. By integrating the findings of the present study with the knowledge from existing literature, we propose a comprehensive model of somatosensory processing that includes both the hierarchical organization within each “major” level and the flow of information between areas. Furthermore, we quantify the structure-receptor and receptor-function couplings by comparing the established RC with the SC and FC of the macaque somatosensory cortex (Fig. 1). The differences among these three modalities provide insights into how structure-function relationships can be modulated by neurotransmitter receptors.

**Figure 1.**
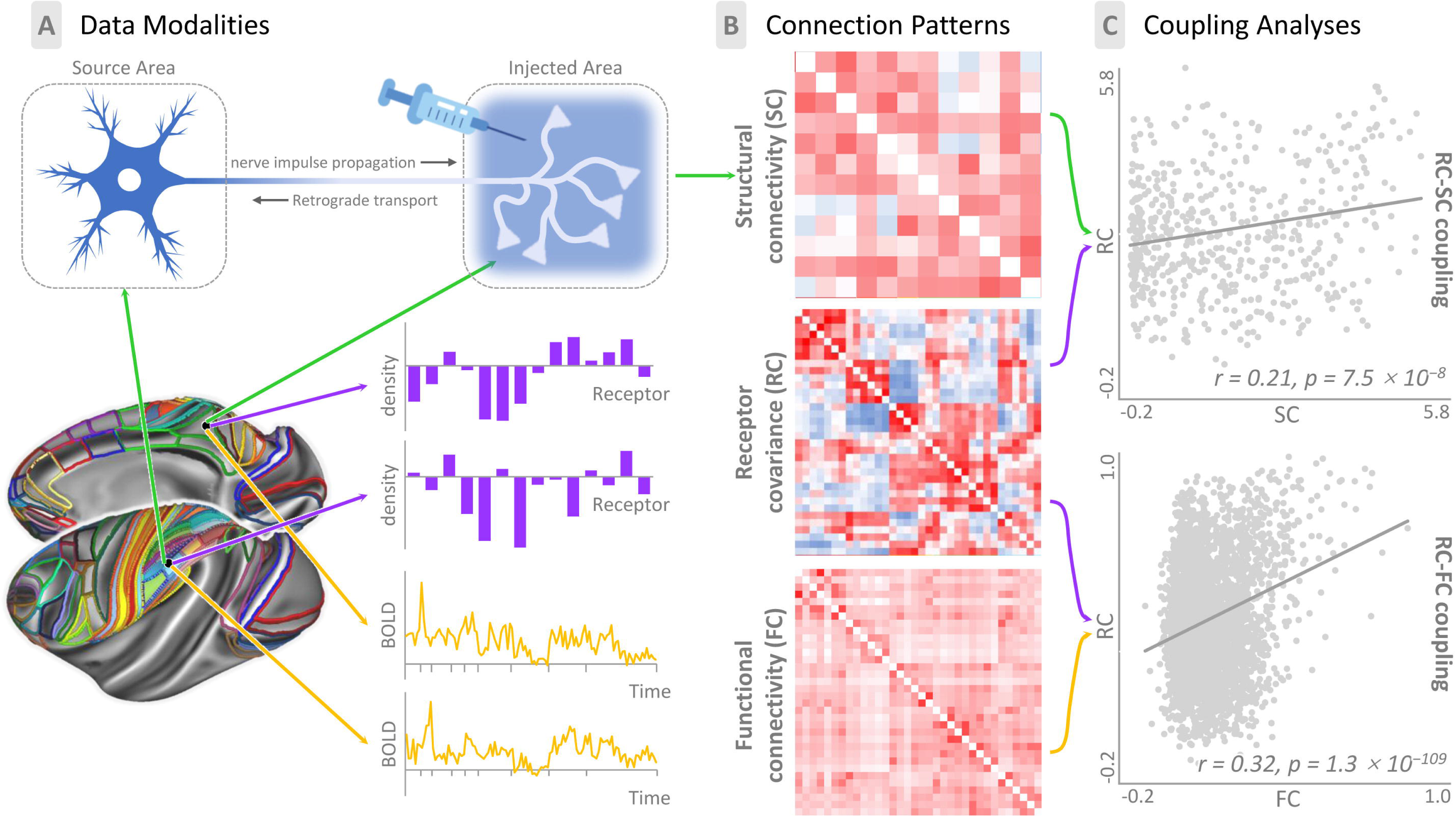
Constructing and comparing the structural connectivity (SC), receptor covariance (RC) and functional connectivity (FC) patterns in the macaque somatosensory cortex. **A:** Extract data used to build the SC (top), RC (middle) and FC (bottom) patterns. Bottom left: The seed areas include all somatosensory-related areas in the parietal lobe, and the target areas include other areas covering the cortical surface except for the temporal lobe. The seed somatosensory areas were divided into five groups based on their hierarchical positions in the somatosensory process or anatomical location: primary somatosensory (SI), secondary somatosensory (SII), intraparietal sulcus (*ips*), inferior parietal (IPL) and superior parietal (SPL). The target areas were first divided based on different brain lobes; within each brain lobe, target areas were grouped together with respect to functional systems based on prior knowledge. Top: A retrograde tracer was injected into each brain area. The tracer was retrogradely transported from axon terminals to the cell bodies of neurons that projected to this area. Middle right: the mean density of each of the 14 receptor types in each defined area was extracted by means of quantitative in vitro receptor autoradiography. Bottom right: the representative activity time course for each defined area was obtained from the resting-state MRI images. **B:** The SC matrix was reconstructed using the fraction of labeled neurons (FLN), which represents the strength of inter-areal SC between each seed area and target area. The RC and FC patterns were reconstructed using representative receptor feature vectors and representative activity time courses, respectively. The statistical similarity between the two areas in RC and FC was evaluated by calculating the Pearson correlation. **C:** The RC-FC and RC-SC couplings were measured by computing the Pearson correlation between the RC and FC or RC and SC vectors, respectively. The RC is positively correlated with FC (Pearson’s *r* = 0.32, p = 1.3 ×10^−109^) and SC (Pearson’s *r* = 0.21, *p* = 7.5 ×10^−8^) patterns.

## Results

To investigate the comprehensive connectivity patterns of the macaque somatosensory cortex, we started by defining the seed and target areas and categorizing them into distinct groups as follows. The seed somatosensory areas were divided into five groups based on their hierarchical positions in the somatosensory process or anatomical location: primary somatosensory (SI), secondary somatosensory (SII), superior parietal (SPL), intraparietal sulcus (*ips*) and inferior parietal (IPL). The target areas were first divided based on different brain lobes; within each brain lobe, target areas were grouped together with respect to functional systems based on prior knowledge. Subsequently, we constructed the RC (Fig. 2), FC (Fig. 3) and SC (Fig. 4) profiles for each somatosensory area using in-vitro receptor autoradiography, resting-state fMRI, and retrograde tracer data, respectively.

**Figure 2.**
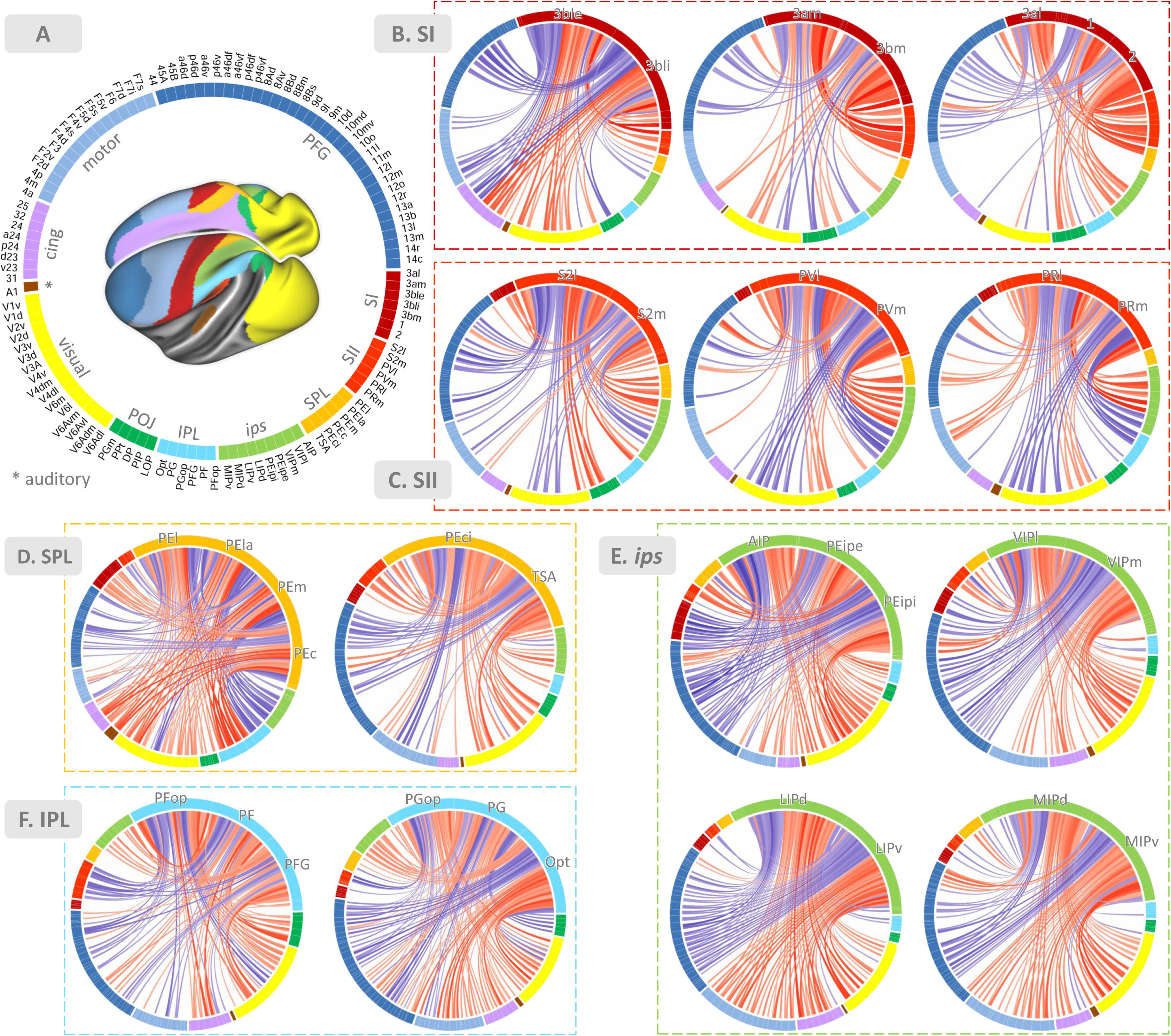
Receptor covariance (RC) patterns of somatosensory-related areas are displayed on the circular plots. The lines connecting two areas represent the covariance strength between the corresponding two brain areas, the thicker the line, the stronger the covariance. The red lines represent positive covariances and the blue lines represent negative covariances. Areas with the same color belong to the same functional or structural subsystem. The color and relative position of each brain area are identical in all circular plots and specified in **A**. **B-F:** The connectograms show the receptor covariance patterns of SI (**B**), SII (**C**), SPL (**D**), *ips* (**E**) and IPL (**F**) areas.

**Figure 3.**
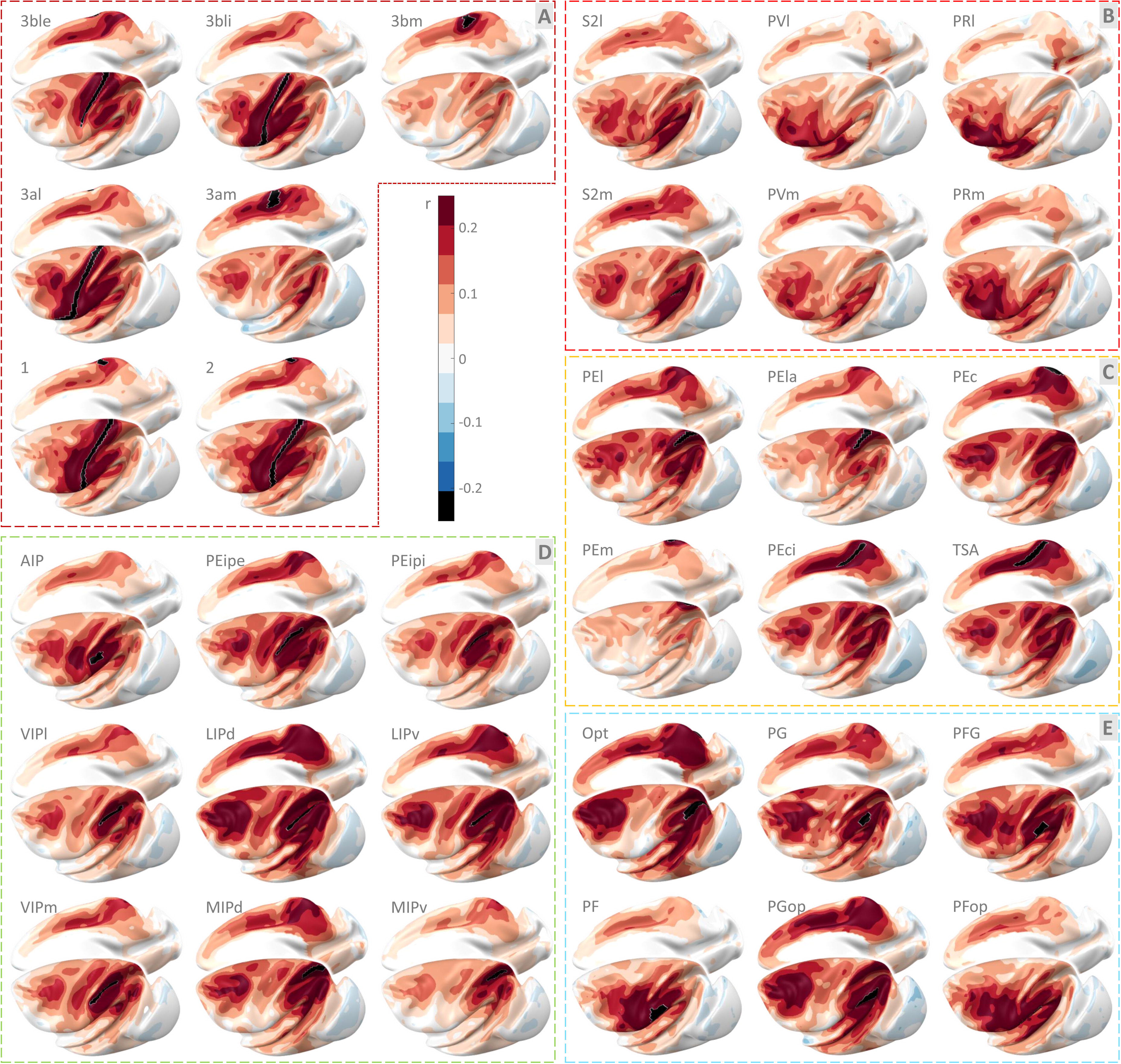
The resting-state functional connectivity patterns of somatosensory-related areas are projected onto the Yerkes19 surface. Colors indicate the strength of Pearson correlation between the timecourse of the seed region (shown in black, summarised by its first principal component) and the timecourse of activity in every other vertex in the cortex.

**Figure 4.**
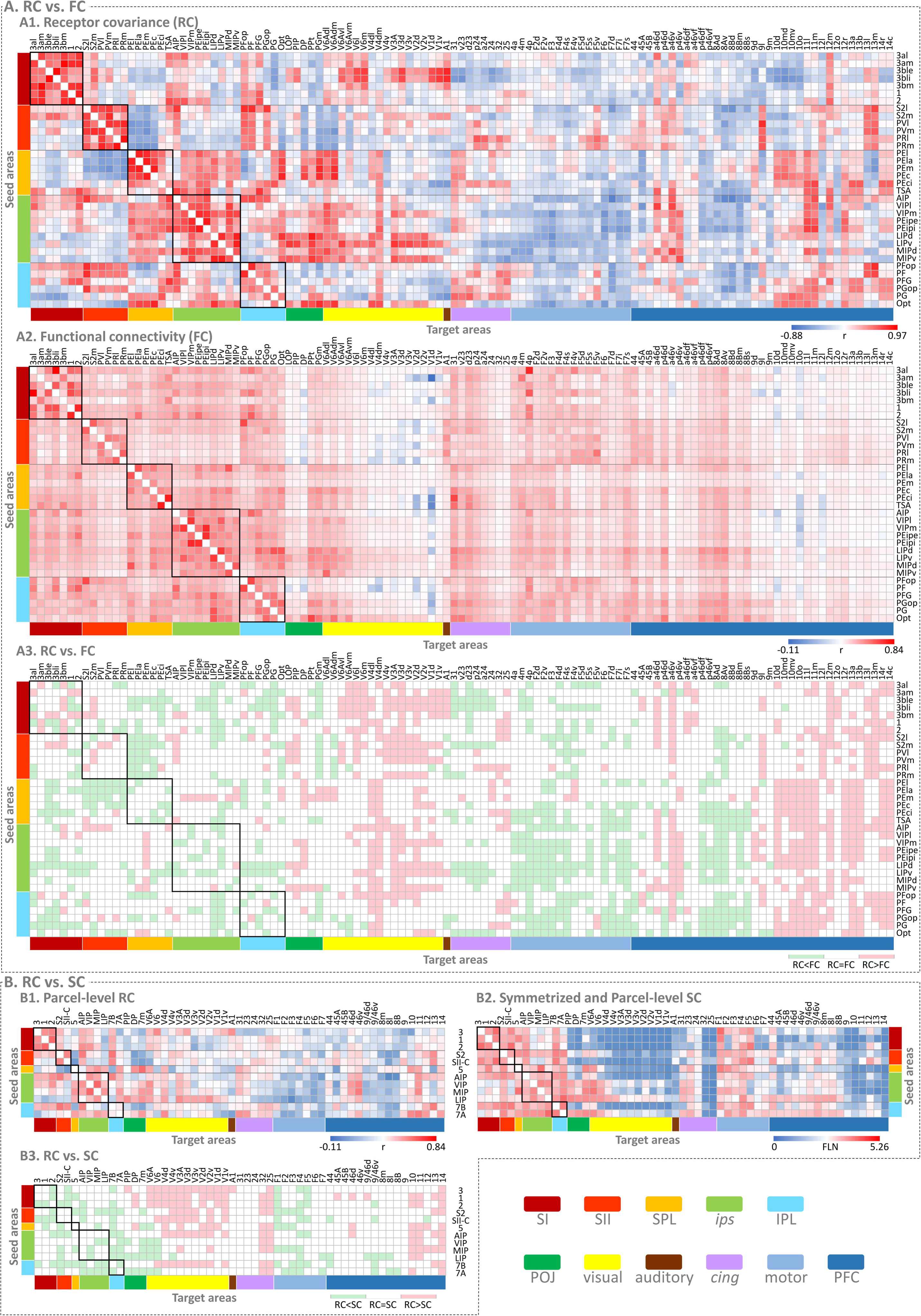
Comparisons between receptor covariance (RC), functional connectivity (FC) and structural connectivity (SC) matrices in macaque somatosensory cortex. **A:** Comparison between RC and FC. The connectivity matrices of RC (**A1**) and FC (**A2**) were constructed based on the MEBRAINS atlas. Each row represents 1 of the 34 seed areas; each column represents 1 of the 115 target areas. The seed areas were divided into five groups based on their hierarchical positions in the somatosensory process or anatomical location: primary somatosensory (SI), secondary somatosensory (SII), intraparietal sulcus (*ips*), inferior parietal (IPL) and superior parietal (SPL). The target areas were first divided based on different brain lobes; within each brain lobe, target areas were grouped concerning functional systems based on prior knowledge. The color bars show the correlation/connectivity strength. The comparison matrix (**A3**) was generated by subtracting the FC matrix from the RC matrix. Before subtraction, the values in both the FC and RC matrices were independently normalized using z-scores. It shows areas of similarity (white) and difference (red/green) in FC and RC between each pair of areas. **B:** Comparison between RC and SC. To compare RC and SC, the original receptor and tracer data were integrated into a common atlas. **B1:** The parcel-level RC matrix. Small regions in the MEBRAINS atlas were merged into larger areas in the common atlas, and their receptor densities were averaged to represent the receptor density of each larger area. **B2:** A symmetrized and parcel-level RC matrix. A symmetrized SC matrix was first obtained by averaging the FLN values from two asymmetric SC matrices, which were constructed based on the Lyon atlas. One of these is an asymmetric 12_target_ ×65_source_ matrix, where somatosensory-related areas serve as the injected areas. The other is an asymmetric 12_source_ × 65_target_ matrix, where somatosensory-related areas serve as source areas (Supplementary Fig. 1). The symmetrized RC matrix was then integrated into the symmetrized and parcel-level RC matrix. In this process, small areas in the Lyon atlas were merged into larger regions in the common atlas, and their FLN values were averaged to represent the FLN values of the larger areas. **B3:** The comparison matrix was generated by subtracting the symmetrized and parcel-level SC matrix from the parcel-level RC matrix. Before subtraction, both matrices were independently normalized using z-scores. The resulting matrix highlights areas of similarity (white) and differences (red/green) in SC and RC between each pair of regions. The color bars indicate FLN weight.

### Receptor covariance (RC) pattern of the somatosensory cortex

A group-average RC pattern for all somatosensory-related areas was constructed across all hemispheres studied (Fig. 2). Visual inspection of the similarity matrix indicated that anatomically adjacent areas share numerous similarities in their RC patterns (Fig. 4A1). Additionally, RC patterns are similar among areas within the same intrinsic network or at the same hierarchical level of brain organization. For example, all SI areas exhibited similar RC patterns, but these patterns differed from those of SII areas (Figs. 2 and 4A1).

#### RC pattern of the SI areas

All SI areas were primarily correlated with each other and showed consistent correlations with caudal SII area S2, the transitional sensory area (TSA), rostral and ventral *ips* areas (AIP, VIP), rostral IPL areas (PFG, PF, PFop), higher visual areas (V6, V6Av), as well as the anterior subdivisions of prefrontal areas 46 and 12m.

Within the SI cortex, areas 3bli and 3ble were distinguished from other SI areas by their unique RC patterns. Compared to other SI areas, 3bli and 3ble exhibited stronger RC with the SII areas S2m and PVm, as well as most of the visual areas. Specifically, they demonstrated the highest RC with the primary visual (V1) and primary auditory (A1) areas among all somatosensory areas (Figs. 2B and 4A1).

#### RC pattern of the SII areas

In addition to their strong intra-areal correlations, SII areas also showed RC with AIP, rostroventral IPL areas (PGop, PF, PFop), and prefrontal areas 9l, 13l and 13m.

Differences in RC patterns also confirmed the segregation of SII. The primary differences between S2 subdivisions (i.e., S2m, S2l) and other areas (i.e., PVm, PVl, PRm, PRl) were detected in their RC with SI areas. S2 subdivisions showed a significantly positive RC with SI areas, whereas the RC between PV/PR subdivisions and SI areas was barely detectable. PR can be distinguished from S2 and PV by its heightened RC with the anterior cingulate, anterior midcingulate areas, as well as ventrolateral premotor area F5 (Fig. 2C).

#### RC pattern of the SPL areas

All SPL areas showed strong RC with the *ips* areas, higher visual areas V4 and V6Ad, frontal pole area 10, and orbitofrontal area 11. Additionally, moderate correlations were observed with orbitofrontal areas 12m, 12r, 13b, and 14r.

The visualization of RC patterns suggested a segregation within SPL. The four subdivisions located on the SPL surface (i.e., PEl, PEla, PEm and PEm) showed notably strong RC with the most caudal area of the entire parietal lobe, such as areas Opt, DP, PPt and PGm. While, the two subdivisions located within the cingulate gyrus (i.e., PEci and TSA) had more consistent RC with rostral and intermediate IPL. Most importantly, TSA can be separated from other SPL areas due to its stronger RC with most of the SI and SII areas (i.e., 3al, 3am, 3bm, 1, 2, S2m and S2l) (Fig. 2D).

#### RC pattern of the ips areas

The *ips* areas exhibited robust RC with areas located within the *ips* itself, as well as in the SPL, posterior IPL and visual cortex. Additionally, they demonstrated moderate RC with prefrontal area 46 and orbitofrontal areas 11l, 11m, 12m and 12r.

While nearly all *ips* areas displayed widespread RC with other brain regions, variations in RC patterns among *ips* areas were noticeable, particularly in their RC with the parietal and occipital lobes. Interestingly, this inter-areal variability in the RC pattern was not random throughout the *ips*, rather, it demonstrated a gradation when moving from rostral to caudal areas. The most anterior area AIP exhibited strong RC with areas located in the SI and SII cortices, rostral and middle IPL and TSA. Intermediate areas PEip and VIP tended to show reliable but moderate RC to the aforementioned targets, but additionally featured consistent RC to the entire SPL, posterior IPL and higher visual areas. The caudal-most areas displayed minimal RC with SI and SII cortices, SPL, anterior IPL and cingulate, but primarily correlated with all visual areas (Fig. 2E).

#### RC pattern of the IPL areas

Similar to areas of the *ips*, IPL areas also exhibited a rostro-caudal gradation in their RC patterns. Rostral areas PF and PFop generally displayed stronger RC with SI and SII areas, dorsal posterior cingulate, posterior midcingulate areas, as well as orbitofrontal areas 13l and 13m. Conversely, the caudal-most area Opt demonstrated stronger connections with *ips*, IPL, and visual areas, as well as with frontopolar area 10, and orbitofrontal area 11 (Fig. 2F).

### Functional connectivity (FC) pattern of the somatosensory cortex

A group-average FC pattern for all somatosensory-related areas was constructed across all studied hemispheres and shown in Figs. 3 and 4A2. Similar to RC patterns, the strongest FC was observed between neighboring areas and within a given intrinsic network.

#### FC pattern of the SI areas

The visualization of FC patterns indicated a clear segregation between lateral (i.e., 3al, 3ble, 3bli, 1, 2) and medial (i.e., 3am, 3bm) subdivisions. In terms of intra-area FC, the lateral areas showed consistent FC with each other, but were barely connected with the medial areas. Moreover, lateral areas presented more prominent FC with widespread brain areas compared to medial areas. All the lateral SI areas showed prominent correlations with rostral and ventral *ips* areas (AIP, VIP, PEip), rostral IPL areas (PFG, PF, PFop) and primary motor area 4p. In contrast, the connections of medial SI areas were primarily confined to areas located on the medial surface of the hemisphere (i.e., TSA, 4m, F3) (Fig. 3A).

#### FC pattern of the SII areas

SII areas also exhibited strong intrinsic FC, with the strongest ones observed between each paired medio-lateral subdivision (i.e., S2m vs. S2l, PVm vs. PVl, PRm vs. PRl). Moreover, SII areas also showed prominent FC with frontal and parietal operculum and moderate FC with widespread brain areas.

Interestingly, the FC pattern shifted from caudal to rostral SII: caudal areas S2m and S2l showed stronger FC with parietal temporal junction and more widespread FC with somatosensory-related areas which are located in SI, SPL, *ips* and IPL. While rostral areas PRm and PRl were predominantly connected to premotor (i.e., F4, F5) and prefrontal (i.e., posterior 46, 8A) areas (Fig. 3B).

#### FC pattern of the SPL areas

Within the SPL, area PEm stood out due to its prominent inter-areal FC with the remaining SPL areas, while exhibiting only limited FC with other areas across the brain. Conversely, other SPL areas displayed a more widespread FC pattern with various distinct areas across the brain. Specifically, most SPL areas exhibited robust and consistent connections with areas situated on the lateral and medial banks of the *ips* (i.e., PEip, LIP, MIP), caudal portion of IPL (i.e., Opt, PG, PGop), as well as the posterior cingulate cortex (i.e., 31, 23d). Moreover, they displayed moderate FC with SI, the *ips* fundus (VIP), parietal-occipital junction (i.e., PGm, V6Adm, V6Adl), primary motor (i.e., 4a, 4p, 4m), premotor (i.e., F2, F3, F7), prefrontal (i.e., p46d, p46df, 8Ad, 8Av) and cingulate cortex (i.e., d23ab, 23c, 24d) (Fig. 3C).

#### FC pattern of the ips areas

In addition to demonstrating robust intra-areal FC, the *ips* areas also exhibited widespread FC with various distinct areas in anterior parietal, SPL, IPL, higher visual, premotor and cingulate cortex.

Visual comparison of the FC patterns revealed a fundamental distinction among all *ips* areas. Generally, FC appeared to be broader in areas closer to the brain surface compared to those situated deeper within the sulcus. Moreover, a variation in FC was appreciable when moving from caudal (e.g., MIP, LIP) to rostral (e.g., AIP, PEip) parts of *ips*. When comparing rostral and caudal areas, it was evident that while caudal regions exhibited more limited FC with SI areas, they demonstrated broader FC with regions spanning the posterior parietal, parieto-occipital junction, primary motor cortex, premotor cortex, and prefrontal cortex (Fig. 3D).

#### FC pattern of the IPL areas

Similar to *ips*, IPL areas also displayed a prominent rostro-caudal gradation in their FC patterns. Rostral areas (i.e., PF, PFG, PFop) generally displayed stronger FC with SI areas 3al, 3bl, 1 and 2, as well as with areas in SII and the anterior *ips*. Conversely, the caudal-part (i.e., Opt, PG, PGop) demonstrated stronger FC with caudal *ips*, SPL, parieto-occipital junction (i.e., PGm, V6Ad, V6Av) and prefrontal areas (Fig. 3E).

### Structural connectivity (SC) patterns of macaque somatosensory cortex

The weights and directions of interareal connections were quantified based on retrograde tracing data, allowing us to create two asymmetric SC matrices: one displays somatosensory-related areas as the injected areas (Supplementary Fig. 1A; 12_injected_ × 65_source_ injected matrix) and provides information concerning the areas that project to somatosensory areas, i.e., the afferents to somatosensory areas. The other matrix shows somatosensory-related areas as the source areas (Supplementary Fig. 1B; 12_source_ × 65_injected_ source matrix) and provides information concerning the areas that receive projections from somatosensory areas, i.e., the efferents from somatosensory areas.

Generally, the connection range of SI areas was limited to a relatively restricted region. Areas 3 and 1 sent more projections than they received, while the number of inputs for area 2 was comparable to its output. SI areas mainly sent projections to other areas within the SI, SII cortex, anterior and lateral *ips* (i.e., AIP and LIP), anterior IPL (7B), primary motor (4), premotor areas (i.e., F4 and F5), and prefrontal area 44. They also received feedback input from most of these areas, except for LIP.

Compared to SI areas, SII areas displayed more extensive SC patterns, with more bidirectional but fewer unidirectional connections than SI areas. SII areas not only received strong input from areas 3, 1, 2, 5, AIP, 7B, 7A, and 4, but also sparse input from areas located in the *ips*, premotor, and prefrontal cortex. Although S2 and the SII complex had a similar number of inputs, S2 showed more output than the SII complex. Specifically, regarding connections with other parietal areas, S2 almost evenly sent projections to all parietal areas, including those located in the medial parietal and parietal-occipital junction, while the SII complex mainly sent output to AIP, LIP, and 7B.

The somatosensory association areas presented more extensive and complex SC patterns than those of the SI and SII areas, especially since they send more projections to other brain areas compared to the former two. In addition, within the somatosensory association cortex, LIP and 7A can be clearly distinguished from other areas since they had particularly rich and strong input from a larger number of other areas across the whole cortex.

### Coupling analyses

#### RC-FC coupling of somatosensory areas

Generally, we found that the RC matrix was significantly correlated with the FC matrix when considering all areas (Fig. 1C; Pearson’s *r* = 0.32, p = 1.3 × 10^−109^). For specifically somatosensory-related areas, the RC-FC couplings ranged from 0.0915 to 0.4893 (Fig. 5A).

**Figure 5.**
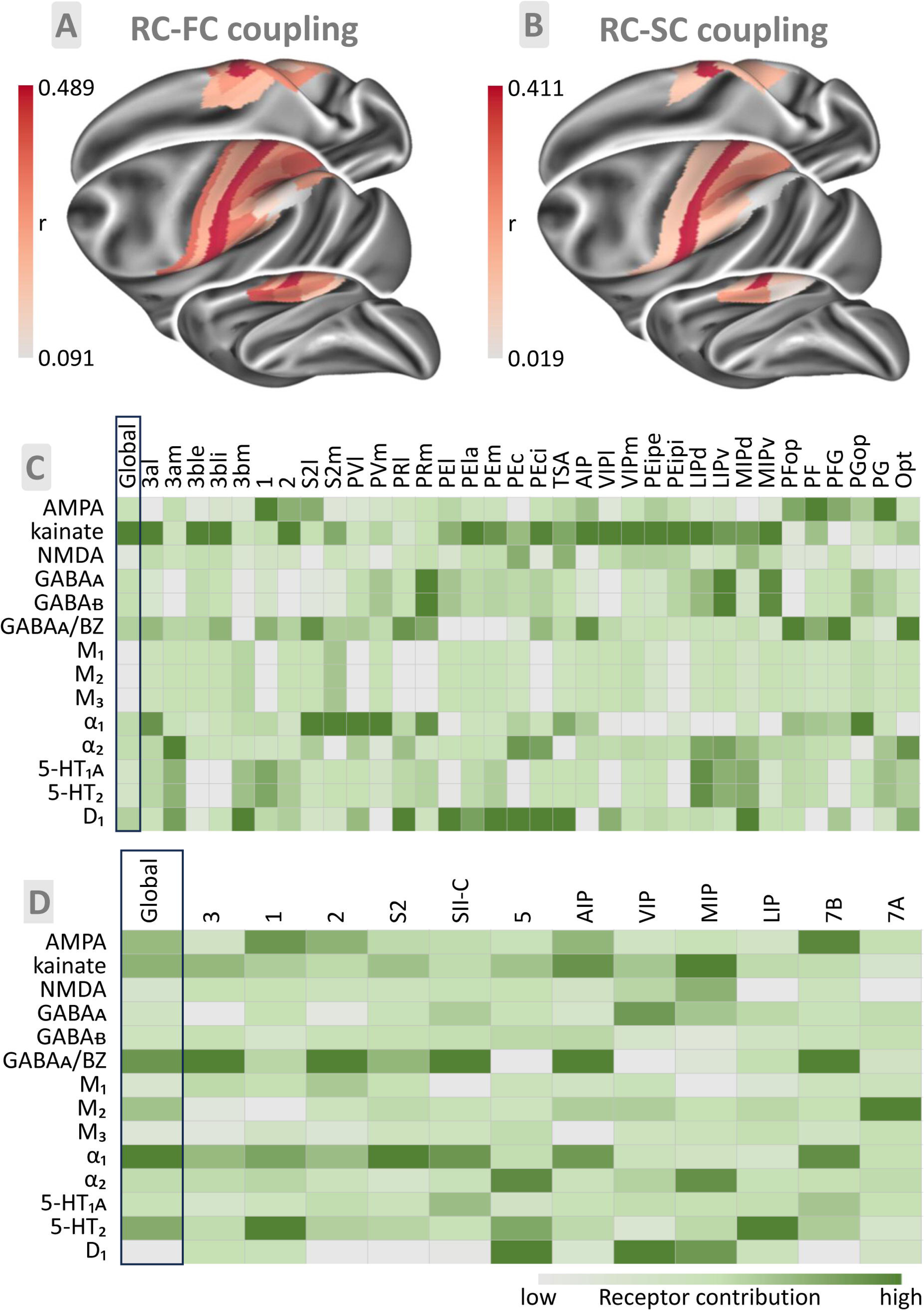
RC-FC and RC-SC couplings of the macaque somatosensory cortex. **A:** RC-FC couplings of all somatosensory areas, ranging from 0.0915 to 0.4893, are displayed on the Yerkes19 surface. **B:** RC-SC couplings of all somatosensory areas, ranging from 0.0190 to 0.4108, are displayed on the Yerkes19 surface. Color bars in (**A**) and (**B**) code for coupling strength of each area, red indicated more similarities between RC-FC or RC-SC, and grey indicated less similarities. **C:** The receptor contribution in RC-FC coupling. The data in each column were relatively independent and represented the contribution of all examined receptors in each brain area. For each column, the color of each entry indicated the contribution of a receptor to the RC-FC coupling (green indicated a higher receptor contribution to the coupling, and grey indicated a lower contribution). **D:** The receptor contribution in RC-SC coupling. For detailed information, see **C**.

The most striking similarities between RC and FC patterns were primarily observed in the strong correlations or connectivity patterns between somatosensory-related areas and their anatomically adjacent areas, or areas at similar hierarchical levels in somatosensory processing. (Fig. 4A3). Despite sharing common features, RC and FC patterns appeared to present more differences than commonalities. The most notable distinction was that, overall, RC exhibited stronger connections and also more negative connections compared to FC. Furthermore, compared to FC, RC demonstrated stronger correlations with visual areas and the anterior/orbital part of the PFC, but weaker correlations with motor, premotor, and posterior PFC regions. In addition, the variability of RC patterns among somatosensory areas was more pronounced than that of FC.

Specifically, the difference between RC and FC patterns can be observed when examining the connections between somatosensory areas and other cortical areas situated in different functional systems. For SI areas, in comparison with FC, RC patterns exhibited stronger correlations with visual areas and areas situated within the rostral principal sulcus, but weaker correlations with motor areas and areas located within the caudal principal sulcus and posterior cingulate cortex. Concerning SII, *ips* and IPL areas, the rostro-caudal segregation was more pronounced in RC patterns compared to FC patterns. In terms of SPL areas, RC patterns displayed stronger correlations with areas of the *ips*, but weaker correlations with SI and SII areas compared to FC patterns.

#### RC-SC coupling of somatosensory areas

To enable the comparison between RC and SC, it was first necessary to integrate the tracer and receptor data into a common atlas to ensure that both matrices had the same number of rows and columns. For this step, we quantified the RC matrix at the parcel level (Fig. 4B1) and symmetrized the SC matrix by averaging the injected and the source SC matrices (Fig. 4B2).

RC was significantly positively correlated with both SC and FC (Fig. 1C). However, the correlation between RC and SC (Fig. 1C; Pearson’s *r* = 0.21, *p* = 7.5 × 10^−8^) was weaker than that between RC and FC (Fig. 1C; Pearson’s *r* = 0.32, *p* = 1.3 × 10^−109^). For specific somatosensory-related areas, the RC-SC couplings ranged from 0.0190 to 0.4108 (Fig. 5B).

In both RC and SC patterns, somatosensory-related areas demonstrated strong correlations or connectivity with their neighboring areas (Fig. 4B). However, unlike the RC pattern, where the strongest correlations were only found between areas within the same hierarchical levels, strong SC patterns were observed between almost all somatosensory areas. In addition, the homogeneity of SC patterns among all somatosensory areas was more pronounced than that of their RC patterns (Figs. 4B1 and 4B2). All somatosensory areas had strong SC with motor and premotor areas as well as prefrontal area 44. This connection pattern also reflected the most significant differences between RC and SC patterns: in terms of the connectivity with areas from other functional systems, somatosensory areas displayed strong SC with primary motor and premotor areas, but strong RC with primary visual and primary auditory areas. Overall, SC exhibited stronger connectivity patterns than RC. For a specific area, the connection strength of its SC patterns varies more widely than that of its RC patterns (Figs. 4B1 and 4B2).

### Receptor contribution analysis

The next question that needs to be addressed is which receptors contribute most to the RC-FC or RC-SC couplings. The matrices in Fig. 5 showed the receptor contribution in RC-FC (Fig. 5C) and RC-SC (Fig. 5D) couplings. The data in each column were independent tested using ‘leave-one-out’ analysis across receptors and represented the contribution of all examined receptors in each brain area. For each column, the color of each entry indicated the contribution of a receptor to the coupling (green indicated a higher receptor contribution to the coupling, and grey indicated a lower contribution).

Overall, the examined 14 receptors influenced RC-FC couplings in a coordinated manner, with no single receptor having a dominant effect. However, kainate, GABAᴀ/BZ, α_1_, α_2_ and D_1_ receptors contributed relatively more to global RC-FC couplings than other receptors. Across brain areas, the contribution of individual receptors varied to RC-FC couplings. For the SI cortex, kainate contributed mostly to the RC-FC couplings of most areas (3al, 3bli, 3ble, and 2), while 5-HT_1A_, 5-HT_2_, and D_1_ receptors also showed high contribution to those of areas 3am, 3bm, 1, and 2. The RC-FC couplings of SII areas are mainly influenced by the GABAᴀ/BZ and α_1_ receptors. The kainate and D_1_ receptors contributed mostly to the RC-FC couplings of all SPL areas. In terms of the *ips* areas, in addition to the kainate receptor contributing to RC-FC coupling in all areas, the GABA_A_, GABA_B_, α_2_, 5-HT_1A_, and 5-HT_2_ receptors also predominantly affect the coupling of caudal *ips* areas. Finally, the AMPA, GABA_A_/BZ, and α_1_ receptors exerted varying degrees of promotion on the FC-RC couplings of the IPL areas.

Generally, the kainate, GABAᴀ/BZ, α_1_, and 5-HT_2_ receptors contributed most to overall RC-SC couplings. Different receptors contributed variably to area-specific RC-SC couplings. Within the SI cortex, the kainate, GABAᴀ/BZ and α_1_ receptors contributed mostly to the RC-FC coupling of area 3, the AMPA, α_1_, and 5-HT_2_ receptors had the greatest impact on that of area 1, and the GABAᴀ/BZ receptor showed the highest contribution to that of area 2. The GABAᴀ/BZ and α_1_ receptors contributed mostly to the RC-SC couplings of SII areas. The RC-SC couplings of SPL are mainly influenced by the α_1_, 5-HT_2_ and D_1_ receptors. Within the *ips*, the AMPA, kainate, GABAᴀ/BZ and α_1_ receptors contributed mostly to the RC-FC coupling of area AIP, the kainate, NMDA, GABAᴀ, α_2_, and D_1_ receptors had different degrees of influence on that of areas VIP and MIP, and the 5-HT_2_ receptor showed the highest contribution to that of area LIP. In terms of the IPL, the RC-SC coupling of the anterior part is predominantly affected by AMPA, GABAᴀ/BZ, and α_1_ receptors, while that of the posterior part is mainly influenced by the M_2_ receptor.

### Cluster Analysis

Multivariate analyses unveiled distinct organizational principles among RC, FC and RC patterns in somatosensory-related areas. For RC, five clusters were identified by k-means clustering analysis. **Cluster 1** included all areas of SI; **Cluster 2** comprised all SII areas along with anterior IPL areas PF and PFop; **Cluster 3** consisted of two medial SPL areas (PEci and TSA), anterior *ips* area AIP, as well as areas in the middle of IPL (PG, PFG, and PGop); **Cluster 4** encompassed four SPL areas situated on the dorso-lateral surface (PEl, PEla, PEm and PEc); and **Cluster 5**, the largest cluster, encompassed the majority of *ips* areas combined with Opt (Fig. 6A).

**Figure 6.**
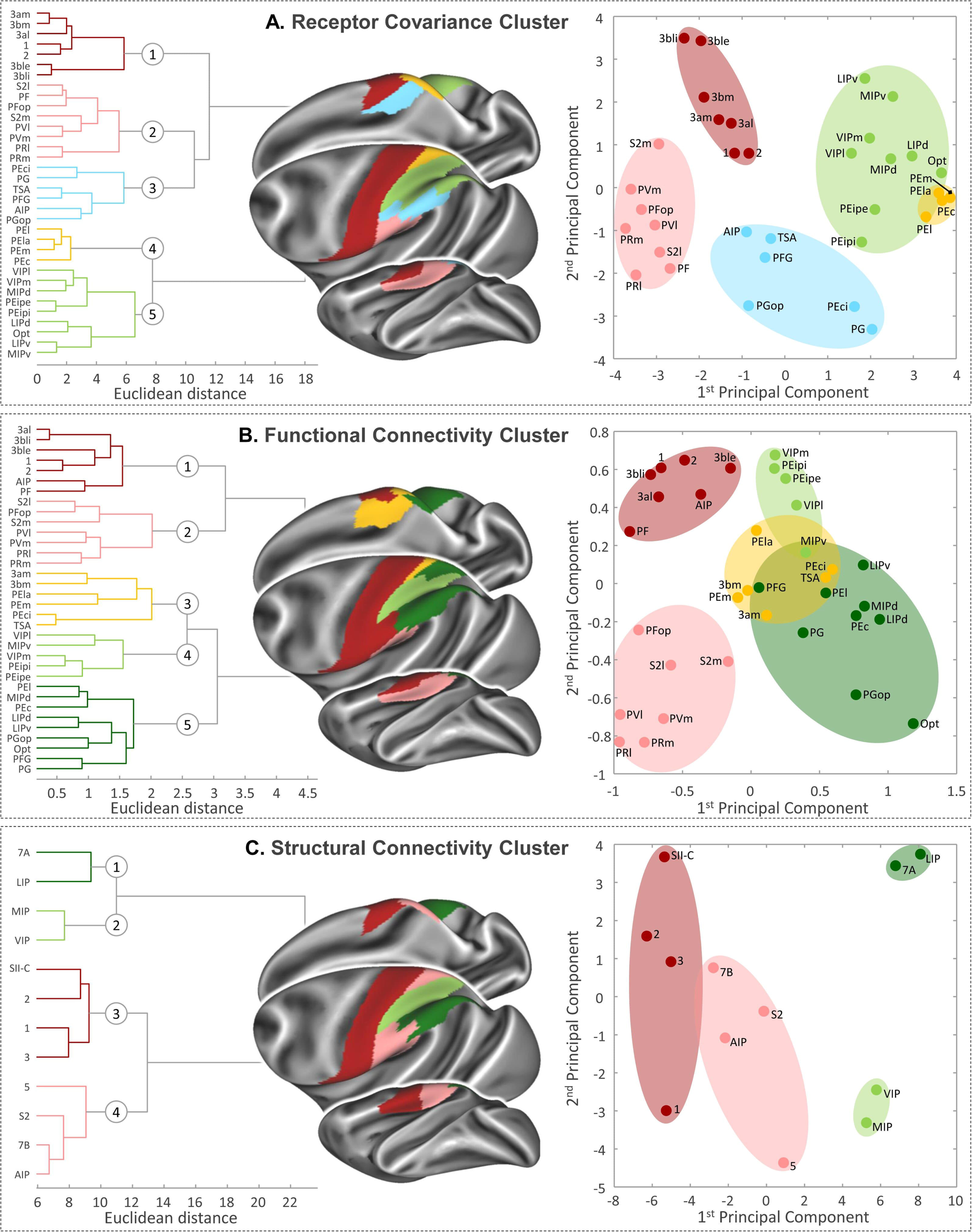
Comparison among RC-driven (**A**), FC-driven (**B**) and SC-driven clustering (**C**) of the macaque monkey somatosensory network. Hierarchical cluster analysis reveals distinct clusters in the RC (**A**), FC (**B**) and SC (**C**) of the somatosensory cortex (left). The detected clusters are displayed on the Yerkes19 surface (middle). Principal Component Analysis (PCA) of the RC (**A**), FC (**B**) and SC (**C**) of somatosensory related areas (right).

The FC can be separated into five clusters as well, **Cluster 1** consisted of the SI areas located on the lateral part (3al, 3ble, 3bli, 1, and 2), along with the anterior part of the *ips* and IPL (AIP and PFop); **Cluster 2** included all SII areas, as well as the rostralmost IPL area PFop; **Cluster 3** comprised two medial SI areas (3am and 3bm) and four SPL areas positioned anteriorly and medially (PEci, TSA, PEm, and PEla); **Cluster 4** contained areas located on the anterior part or near the depth of the *ips* (VIPl, VIPm, PEipe, PEipi, and MIPv); and as the largest cluster, **Cluster 5** was composed of nine areas situated on the caudal part of SPL, *ips*, and IPL (Fig. 6B).

The SC was separated into four clusters based on the k-means analysis, **Cluster 1** consisted of the SI areas located on the lateral part of the hemisphere (3, 1, and 2) along with the SII complex areas; **Cluster 2** included areas S2, 5, AIP and 7B; **Cluster 3** comprised MIP and VIP; **Cluster 4** contained LIP and 7A (Fig. 6C).

Across all three modalities, SI areas, particularly the lateral part, consistently formed a distinct cluster, while SII areas clustered together with the most anterior IPL. The clustering of association areas is more complex, varying between two or three clusters depending on the modality. Regarding the distances between these clusters, the SC-based cluster exhibited the greatest separation, the FC-based cluster had the shortest distance, and the RC-based clusters fell in between.

### Three-stream model of somatosensory processing

By combining the findings of the present study with knowledge from existing literature (See the first paragraph of the Discussion for the pertinent references), we propose a model of somatosensory processing streams in the macaque cortex (Fig. 7). Area 3bl serves as the initial cortical stage, receiving somatosensory signals from the thalamus and projecting horizontally to areas 3al, 1, and 2, thereby establishing a hierarchical structure for early sensory processing.

**Figure 7.**
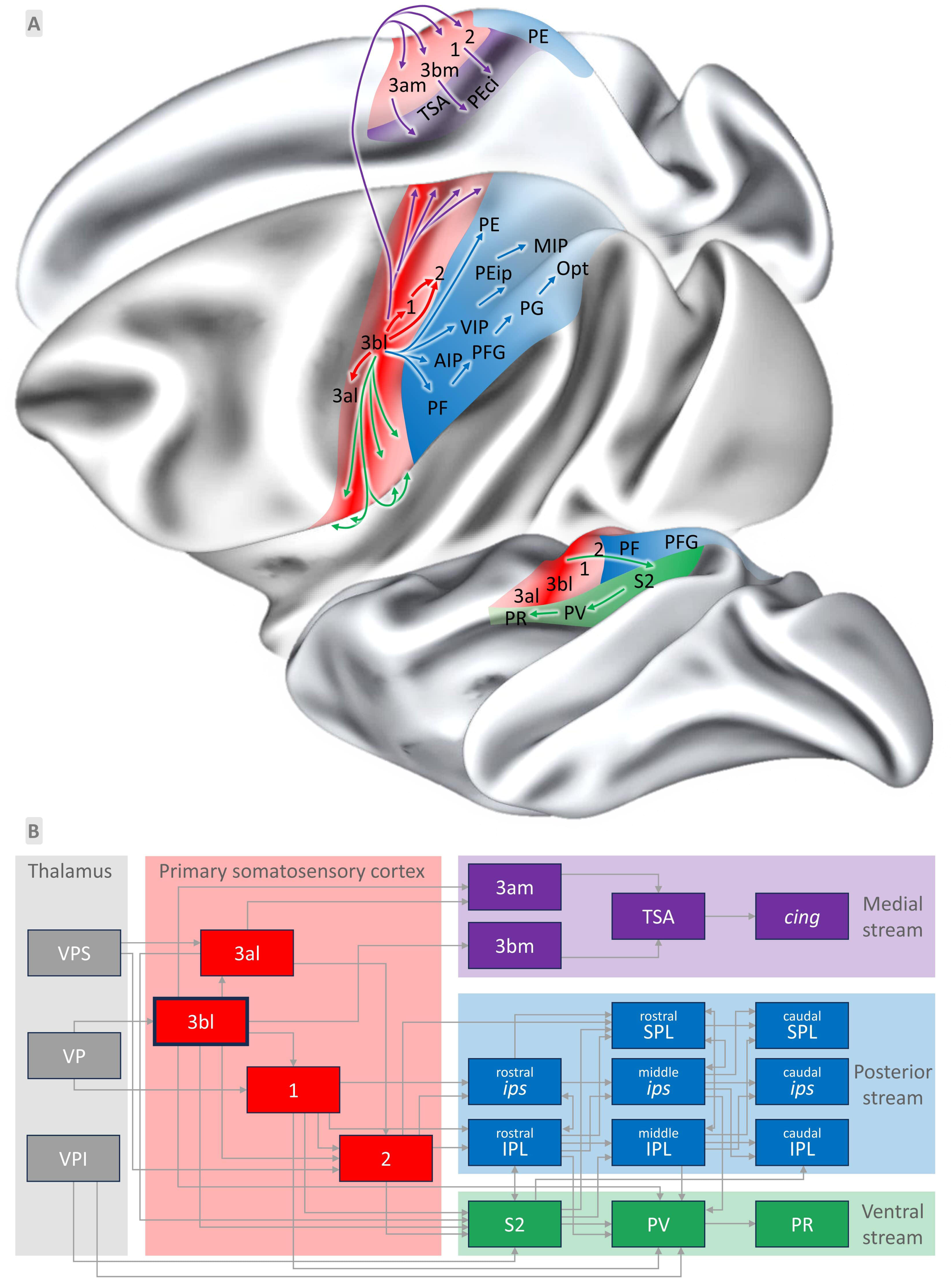
Schematic diagram of the somatosensory transmission pathway and hierarchical organization proposed in the present study. **A**: The schematic diagram illustrates the basic components of the somatosensory pathways in the macaque monkey cortex. Arrows of different colors represent distinct information flows. Area 3bl initially receives somatosensory signals from the thalamus and projects to areas 3al, 1, and 2 within SI (red zone). From there, the lateral SI areas project in three main directions: ventrally to the SII complex (green zone), medially to the medial SI and TSA (violet zone), and posteriorly to somatosensory association areas along the parietal lobe (blue zone). Within each zone, the color purity reflects the hierarchical organization, with higher purity indicating a lower hierarchical level and lower purity indicating a higher level. **B**: A proposed hierarchy of somatosensory-related areas in the macaque monkey cortex. The information flow in **B** aligns with that in **A** but offers a more detailed overview of sensory information transmission for each brain area or region. Hierarchical levels of the areas or regions increase progressively from left to right.

From these lateral SI areas, somatosensory information propagates along three major pathways:

- The medial stream projects to the medial wall of the hemisphere. Somatosensory information is first directed to the medial SI (i.e., areas 3am and 3bm) and then extends to the transitional somatosensory area (TSA) before reaching the cingulate cortex and medial parietal cortex.
- The posterior stream projects to the association areas in the posterior parietal lobe. The signals are first projected to the anterior portion of the *ips* and IPL (i.e., AIP, PF, and PFop) and then transmitted posteriorly along the parietal lobe.
- The ventral stream projects to the secondary somatosensory complex (SII). Specifically, somatosensory signals are first projected to S2 and then extended anteriorly along the caudal-rostral axis to PV and PR.

## Discussion

In this study, we focused on the macaque somatosensory cortex, reconstructing three distinct types of covariance/connectivity patterns: *receptor covariance*, *resting-state functional connectivity*, and *tract-tracing-based structural connectivity*. By combining the key differences in multimodal connectivity patterns detected in the present study with knowledge from existing literature (Felleman and Van Essen, 1991; Kaas, 2012; Rossi-Pool et al., 2021; Saadon-Grosman et al., 2020; Yang et al., 2021), we detect the hierarchical relationships among somatosensory-related areas, and further propose a model of somatosensory processing streams (Fig. 7). Moreover, our findings demonstrate that the RC pattern is closely linked with SC and FC, yet it also exhibits unique characteristics potentially reflect the somatosensory processing hierarchy.

### Hierarchical organization in somatosensory cortex

In the present study, we provide strong evidence from receptor covariance and multimodal covariance patterns to support the hierarchical organization of the somatosensory cortex. Consistent with previous findings (Iwamura, 1998; Kaas, 1993; Rossi-Pool et al., 2021; Saadon-Grosman et al., 2020), our results indicate that the cortical hierarchy for somatosensory processing begins in the SI, progresses to the SII, and continues through the association areas in the posterior parietal cortex. More importantly, we also identified clear hierarchies within SI, SII, and association cortices.

Within SI, area 3bl occupies the lowest hierarchical level in the somatosensory processing. As a prototypical SI area, area 3bl exhibited significantly stronger RC with primary and early visual and auditory areas but relatively weaker RC with association areas compared to other SI components. This aligns with previous evidence showing that area 3b displays faster timescales than other SI areas (Rossi-Pool et al., 2021), and that the receptive fields of area 3b neurons are smaller, simpler and more defined than those in other SI areas (Burton and Fabri, 1995; Iwamura et al., 1983; Iwamura et al., 1993). In terms of the remaining SI areas, area 3a is located in a lower position than area 1 in the somatosensory hierarchy, as it receives direct input from the thalamus and is primarily involved in processing basic proprioceptive information (Darian-Smith et al., 1993; Huffman and Krubitzer, 2001; Krubitzer et al., 2004).

In contrast, area 1 mainly receives projections from areas 3b and 3a, and it handles more complex aspects of sensory information (Burton and Fabri, 1995; Iwamura et al., 1983). Area 2 is involved in a higher level of information processing than area 1, as it displays a rougher somatotopic organization and exhibits much slower timescales (Murray et al., 2014; Pons et al., 1985; Rossi-Pool et al., 2021). Although numerous studies demonstrate a hierarchical relationship among these areas, this is not evident in the multimodal connectivity patterns. In addition, medial areas (i.e., 3am and 3bm) can be differentiated from lateral areas by their relatively weaker RC with other somatosensory-related areas, which implies the medial areas are involved in a higher level of information processing than their lateral counterparts (Niu et al., 2024).

In SII, the hierarchical position gradually increased from S2 to PV to PR. S2 is notably distinct from other SII areas due to its strong SC, RC, and FC with SI areas. Moreover, S2’s connectivity patterns across all three modalities closely resemble those of the SI area. In contrast, PR stands apart from both S2 and PV, characterized by stronger RC with cingulate and premotor areas. In other words, as we move from caudal to rostral areas, connections with SI areas diminish, while connections with association and motor areas become stronger. These findings suggest that somatosensory features are most prominent in S2 and gradually diminish along the S2-PV-PR axis, indicating a possible processing stream that begins in S2 and extends anteriorly. Thus, S2 may occupy an intermediate hierarchical position between SI and other areas within SII in sensory information processing. Although PR is classified as part of SII, it may functionally interface between sensory and motor systems. This hypothesis aligns well with theoretical models that have received experimental support from previous studies (Fitzgerald et al., 2004; Kaas, 1993; Saadon-Grosman et al., 2020).

In comparison to the SI and SII regions, the association areas in the posterior parietal cortex are involved in higher-level information processing, as they show stronger SC, RC, and FC with other association areas. Among these areas, the anterior parts of the *ips* (AIP and VIP) and the IPL (PF and PFop), as well as the medial area TSA, occupy a lower hierarchical position, closer to those of the SI and SII areas. This is due to their stronger SC, RC, and FC with the SI and SII regions. Furthermore, their connectivity patterns more closely resemble those of SI and SII, indicating that their connection patterns exhibit more somatosensory features compared to other association areas.

### Three-stream model of somatosensory processing

Integrating our findings on the hierarchical relationships among somatosensory-related areas with previously published evidence (Arienzo et al., 2006; Mazzola et al., 2006; Saadon-Grosman et al., 2020), we propose a model of somatosensory processing streams in the macaque brain (Fig. 7). In this model, we identify three distinct pathways of information flow from early somatosensory areas to association areas.

Medially, sensory information is transmitted via the medial SI and TSA to the cingulate and medial parietal cortices. Previous studies have provided evidence of somatosensory hierarchies in the medial surface (Kaas, 2012; Morecraft et al., 2004; Niu et al., 2024; Saadon-Grosman et al., 2020). The medial SI areas have larger receptive fields than lateral SI, indicating a higher hierarchical role in the processing stream (Kaas, 2012). As a transitional area, TSA is primarily associated with somatosensory-motor regions and has limited connections with association areas (Morecraft et al., 2004). Our observations indicate that connections with lateral SI areas progressively weaken from the medial end of SI toward the cingulate and medial parietal cortices. This pathway likely plays a role in integrating somatosensory and proprioceptive signals, essential for body awareness, fine motor control, and the modulation of bodily perception in relation to cognitive and emotional states (Laurienti et al., 2003; Morecraft et al., 2004; Saadon-Grosman et al., 2020).

Posteriorly, the signal first reaches the areas closest to the lateral SI, i.e., the frontmost part of the *ips* and IPL, and then spreads across the entire posterior parietal lobe. We observed a progressive decrease in connections with the lateral SI along the rostro-caudal axis in both the *ips* and IPL. This is consistent with previous evidence demonstrating the hierarchical organization of the posterior parietal region (Felleman and Van Essen, 1991; Robinson and Burton, 1980; Rossi-Pool et al., 2021). This pathway is associated with higher-order spatial processing, sensorimotor coordination, and the transformation of sensory input into goal-directed actions, making it crucial for tasks such as grasping and tool use (Caminiti et al., 1991; Fattori et al., 2010; Galletti et al., 1997).Ventrally, the processing stream originates in S2, which occupies a lower hierarchical position within the SII complex. This pathway then continues anteriorly along the caudal-rostral axis to PV and PR (Fitzgerald et al., 2004; Romo et al., 2002).As move from S2 to PR, the connectivity with the SI region decreases, while the connections with other brain areas, particularly the prefrontal cortex, become more extensive. This stream plays a crucial role in higher-order tactile processing, such as texture discrimination and sensorimotor integration (Hinkley et al., 2007; Krubitzer et al., 1995). Furthermore, the PR area is strongly linked to motor processing, helping to relay processed somatosensory information to regions involved in object recognition and multisensory integration (Disbrow et al., 2003; Disbrow et al., 2000).

It’s worth noting that while the hierarchy appears to be highly correlated with information flow across cortical areas, it does not directly correspond to the direction of information transmission during somatosensory processing for specific stimulation. Firstly, information transmission is bidirectional rather than one-way; it does not simply flow from lower to higher levels. For example, area 3b sends signals to area 1 while also receiving parallel feedback inputs from area 1 (Markov et al., 2013; Markov et al., 2014a). Secondly, our model suggests that sensory information originates in area 3b and flows through areas 3a, 1, and 2 to reach SII and association areas. However, connectivity is largely parallel and not serial (Vezoli et al., 2021). For instance, area 3b can send signals directly to areas 2 and SII without passing through area 1 (Baldwin et al., 2018; Krubitzer and Kaas, 1990). Furthermore, the three streams are interconnected, with nodes in different streams also transmitting information to one another (Kaas, 2012; Krubitzer and Kaas, 1990). Overall, the hierarchy reflects the inherent characteristics of each area, giving us insights into which stage of information processing each area primarily occupies. This hierarchy appears to be independent of any specific function or task being studied. Two cortical areas at the same hierarchical level do not necessarily serve the same functions or play the same roles in a task. For example, areas 3b, V1, and A1 share similar hierarchical positions but have sharply distinct functions. Their similar position in the hierarchy simply indicates that each is situated at the first stage of its respective sensory stream.

### Relationship among RC, SC and FC

#### Relationship between RC and SC

In general, areas with anatomical connections tend to exhibit similar receptor fingerprints. This is evident when comparing the RC and SC of SI areas, where SI areas exhibited strong RC with other SI areas, S2, AIP, and anterior IPL. These areas are known to be structurally connected to SI areas based on the SC patterns observed in the present study and previous publications (Burton and Fabri, 1995; Darian-Smith et al., 1993). However, despite the absence of known direct anatomical connections, SI areas also exhibited similarities in RC with most visual areas, primary auditory area, anterior subdivisions of area 46, and prefrontal area 12m. In addition, we also observed strong SC between SI and motor (4) and premotor (F2, F3, F4 and F5) areas, yet these connections were not reflected in their RC. These observations indicate that RC and SC do not have one-to-one correspondence. When cortical areas show strong SC but lack RC, it indicates the presence of significant anatomical connections (such as axonal pathways or white matter tracts) between these regions. However, the synaptic input they receive may not be closely linked to similar sensory receptor activity. These areas may be involved in different sensory or cognitive functions. On the other hand, the absence of direct SC does not necessarily limit RC similarities. Strong RC without SC could be due to polysynaptic connectivity or shared functional networks. Alternatively, it might reflect a similar computational function or hierarchical role within a different network.

As a further example, our results indicated that S2 areas have strong RC with PV, PR, all SI areas of the anterior parietal cortex, as well as the of anterior portions of the *ips* (i.e., AIP) and IPL (i.e., PF, PFop). Structurally, area S2 receives direct projections from areas 3a, 3b, 1, and 2, and to primarily project to PV, PR, and to the anterior portion of posterior parietal cortex (Disbrow et al., 2003). This suggests that RC reflects the information transmission trajectory to some extent, even if it cannot explicitly indicate the directionality of information transmission.

Beyond the similarities mentioned above, RC and SC exhibited more pronounced inconsistencies. First of all, the SC strength is generally higher than that of RC. For pairs of areas with strong connections in both RC and SC, the range of their SC values is over an order of magnitude greater than that of their RC values. Although the RC pattern in the somatosensory areas is weaker than the SC, it is also more extensive. For instance, the SC patterns of the SI areas are highly concentrated, with strong SC connections primarily found with nearby areas and with motor (4) and premotor (F2, F3, F4, and F5) areas. In contrast, RC patterns of the SI areas are more distributed, spanning multiple functional systems beyond anatomical immediate surrounding regions. These inconsistencies arise primarily from the distinct biological natures of RC and SC. The SC patterns involve direct physical pathways, like axon bundles that link various brain areas and are typically stronger to maintain a constant and robust flow of neural signals. In contrast, the RC represents how consistently brain areas respond to neurotransmitters and modulatory functions. RC can be more diffuse and weaker than SC, as it supports flexible and widespread modulatory effects.

Furthermore, we found that the heterogeneity of RC patterns among somatosensory areas is more pronounced than that of SC. This variability in RC patterns across different areas may be related to their positions at different hierarchical levels of somatosensory processing. The prototypical SI area 3bl (Zilles and Palomero-Gallagher, 2020) exhibits distinct RC patterns compared to other SI areas. Subdivisions of 3bl demonstrated significant RC with the primary and secondary visual areas, as well as primary auditory area. In contrast, although subdivisions of areas 3a, 1, and 2 exhibited extensive RC with the anterior IPS, anterior IPL, and higher visual areas V6, V6Av, and V6Ad, there was minimal discernible RC between them and early sensory areas (i.e., V1, V2, and A1). We propose that the variability in RC patterns between area 3bl and other SI areas can be attributed to their different hierarchical levels (Burt et al., 2018; Rossi-Pool et al., 2021). In other words, RC patterns not only reflect direct anatomical connections but can also provide insights into the information processing level within a specific functional system (Froudist-Walsh et al., 2023; Goulas et al., 2021; Rapan et al., 2022). The brain areas with similar multi-receptor balances tend to have the potential to be responsive to similar neurotransmitters or signaling molecules (Palomero-Gallagher et al., 2015). Consequently, they are likely to play similar roles within their respective functional systems. Area 3b, primary visual and primary auditory areas exhibit significant similarity in their receptor fingerprints that are relatively unique with extremely high M_2_ densities (Niu et al., 2024; Rapan et al., 2022; Zilles and Palomero-Gallagher, 2017a), reflecting the importance of the cholinergic system in the modulation of thalamocortical input and improving the signal to noise ratio in cortical processing of sensory stimuli (Herrero et al., 2017; Miyawaki et al., 2023; Zhang and Burger, 2024). Conversely, the receptor fingerprints of association areas are more balanced than those of primary sensory areas, with a higher receptor density per neuron, reflecting their more intensive information exchange (Froudist-Walsh et al., 2023; Goulas et al., 2021; Zilles and Palomero-Gallagher, 2017b).

#### Relationship between RC and FC

Recent studies report that neurotransmitter receptor distributions are systematically aligned with FC at the global level (Froudist-Walsh et al., 2023; Hansen et al., 2022). Our findings support this, as RC and FC patterns exhibit significant correlations in the somatosensory areas, and the cluster analyses of RC and FC exhibited comparable somatosensory grouping. This suggests that both modalities contribute to cortical organizing principles of the somatosensory cortex.

Despite these apparent similarities, we also observed that RC and FC represent different biological foundations for macroscopic connections in the brain. First, the RC pattern of the somatosensory areas is much stronger than their FC pattern in general. This is because the RC pattern provides information about the underlying chemoarchitectonic similarities of different brain areas (Hansen et al., 2022; Palomero-Gallagher et al., 2015). This inherent biological similarity does not change significantly over a short period of time, offering a physical substrate that can support information transmission even in the absence of ongoing neural activity or functional changes (Citri and Malenka, 2008; Kasai et al., 2010). In contrast, resting-state MRI-based FC is represented by the correlation between the BOLD signal time series of different brain areas (Bijsterbosch et al., 2017). It reflects the synchronized neural activity observed between brain areas during rest (Smith et al., 2013), which may not always directly correlate with the underlying architectonic synchronicity (Greicius et al., 2009). Resting-state FC would emerge as a result of ongoing neural activity and can exhibit more variability over time (Bijsterbosch et al., 2017; Smith et al., 2013).

Another key difference is that RC patterns exhibit greater variability than FC. This likely reflects their distinct representations of the foundational organizational principles of the brain. As mentioned previously, the observed variability in RC patterns may reflect differences in the hierarchical level of the examined somatosensory areas. Conversely, the FC reflects the degree to which different brain areas exhibit synchronized or coordinated activity over time (Bijsterbosch et al., 2017; Smith et al., 2013). It can help identify areas that belong to the same functional systems or are activated by the same tasks (Yeo et al., 2011). Therefore, the examined areas displayed similar FC patterns since they are all involved in somatosensory activities (Thomas et al., 2021; Wang et al., 2013).

This observation aligns with our findings: most somatosensory areas exhibit consistent FC with several brain regions, including the PPt, PGm, V6Ad, posterior cingulate, and motor areas such as 4, F2, and F3, as well as the areas located at the caudal portions of the principal sulcus, such as p46 and 8A. Most of these areas exhibit somatosensory-related activity, such as PPt, PGm, V6Ad, 31, 23, and 24d (Cléry et al., 2020; Morecraft et al., 2004; Passarelli et al., 2018; Saadon-Grosman et al., 2020). Cells in areas PGm and V6Ad have been shown to respond to somatosensory stimuli (Gamberini et al., 2018; Gamberini et al., 2011; Passarelli et al., 2018). Previous studies have shown that the posterior and midcingulate (such as 23 and 24d) and precuneus cortex (such as areas 31 and PGm) receive major input from posterior parietal areas (Baleydier and Mauguiere, 1987; Passarelli et al., 2018) involved in processing somatosensory stimuli relevant for reaching and grasping movements (Galletti et al., 2022; Morecraft et al., 1993). Interestingly, despite being the primary motor area, area 4 is often grouped with primary somatosensory areas in functional connectivity studies (Hutchison et al., 2011; Yeo et al., 2011). This reflects the close interplay between primary motor and somatosensory areas in executing motor actions based on sensory input, with primary somatosensory area processing sensory information (Delhaye et al., 2018; Iwamura, 1998; Rossi-Pool et al., 2021) and primary motor area translating this input into motor commands (Darian-Smith et al., 1993; Kurata, 1994; Morel et al., 2005).

Our analysis revealed a significant difference between their RC and FC patterns, with most somatosensory areas exhibiting strong positive correlations with motor areas in the FC pattern, but displaying either negative or weak positive correlations with motor areas in the RC pattern. This is because, although somatosensory areas are deeply functionally integrated with motor areas, their biological features and mechanisms are fundamentally different, leading to distinct roles in their respective functional systems (Delhaye et al., 2018; Kurata, 1994). Primary somatosensory area 3b is characterized by an extremely dense population of small granular cells throughout layers II–IV, along with high densities of M_2_ receptors across the cortex (Niu et al., 2024; Zilles and Palomero-Gallagher, 2020). It is primarily involved in receiving somatosensory input from the thalamus and transporting the information to other somatosensory areas for further processing and analysis (Burton and Fabri, 1995; Darian-Smith et al., 1993). In contrast, the primary motor cortex (area 4) features unusually large pyramidal cells known as Betz cells in sublayer Vb, along with a significantly smaller receptor fingerprint (Rapan et al., 2021), and is responsible for generating voluntary movements and executing motor commands initiated by higher brain regions (Salinas and Romo, 1998). It receives input mainly from pre- and supplementary motor areas, somatosensory cortices, and the thalamus, and sends outputs to the intermediate cerebellum and corticospinal tract (Darian-Smith et al., 1993).

## Conclusion

In summary, the present study demonstrated key differences across somatosensory areas at different hierarchical levels in terms of their multimodal covariance patterns and proposed a comprehensive model of somatosensory processing streams in the macaque brain. In this model, area 3b1 serves as the first cortical stage for somatosensory signals in the primate brain. It then projects horizontally to areas 3a1, 1, and 2, which represent the next level in the processing pathway. From these lateral SI areas, signals propagate in three directions: (1) ventrally to the SII complex along the caudal-rostral axis, (2) medially to the medial SI and TSA, and (3) posteriorly to somatosensory association areas in the parietal lobe.

Furthermore, this study enhances our understanding of brain connectivity across multiple modalities and establishes links among the structural, chemoarchitectonic, and functional organization of the macaque somatosensory cortex. Our findings provide insights into the role of RC in uncovering the fundamental organizational principles of the brain. Compared to SC and FC, RC patterns are more sensitive to hierarchical relationships among brain areas, offering a novel perspective on the multiscale integration of brain structure and function.

## Materials and methods

### Data provenance

To establish the RC, FC, and SC of the macaque somatosensory cortex, we exploited 3 independent publicly available data sets for: i) multiple receptor densities, ii) in vivo MRI data, and iii) retrograde tract-tracing data, respectively (Fig. 1). The RC was generated using multiple receptor densities from previously published studies by the Palomero-Gallagher group (Impieri et al., 2019; Niu et al., 2020; Niu et al., 2024; Niu et al., 2021; Palomero-Gallagher et al., 2013; Rapan et al., 2021; Rapan et al., 2023; Rapan et al., 2022), available via the MEBRAINS Multilevel Macaque Brain Atlas (https://search.kg.ebrains.eu/instances/Project/e39a0407-a98a-480e-9c63-4a2225ddfbe4).

The FC was generated using MRI/fMRI data from the openly available dataset PRIME-DE (Milham et al., 2018a) (http://fcon_1000.projects.nitrc.org/indi/indiPRIME.html). The SC was generated using retrograde track tracing data from the Kennedy lab (Markov et al., 2013; Markov et al., 2014a; Markov et al., 2014b) (https://core-nets.org).

### Definition of regions of interest

To reconstruct the RC, FC, and SC patterns of each somatosensory-related area, we first defined the regions of interest (ROIs). The seed areas refer to the somatosensory-related areas, i.e., SI, SII and the somatosensory association areas located in the SPL, ips, and IPL. In addition to these areas, the target areas for tracer injections were widely distributed across the cortex except those located in the temporal lobe.

The ROIs in the RC and FC analyses differed in extent and terminology from those used in the SC analysis because anatomical measurements of receptor densities and white matter connectivity were analyzed using different atlases. In the context of the RC and FC analyses, the ROIs were defined using the MEBRAINS Multilevel Macaque Brain Atlas (Fig. 1A), whereas the ROIs for the SC analysis were defined based on the Lyon atlas (optimization of the Markov-132 atlas) parcellated by the Kennedy group (Fig. 1A). In both atlases, most areas of the temporal lobe remain unquantified.

To facilitate the description and interpretation of the results, the seed and target areas were summarized into different groups as follows. The seed areas were divided into five groups based on their hierarchical positions in the somatosensory process or anatomical location: primary somatosensory (SI), secondary somatosensory (SII), superior parietal (SPL), intraparietal sulcus (*ips*) and inferior parietal (IPL). The target areas were first divided based on different brain lobes; within each brain lobe, target areas were grouped together with respect to functional systems based on prior knowledge. Detailed information regarding the ROIs can be found in Table S1.

### Receptor covariance (RC) patterns of macaque somatosensory cortex

The quantitative in-vitro receptor autoradiography data used for RC reconstruction was obtained from previously published papers and datasets (Impieri et al., 2019; Niu et al., 2020; Niu et al., 2024; Niu et al., 2021; Palomero-Gallagher et al., 2013; Rapan et al., 2021; Rapan et al., 2023; Rapan et al., 2022) (https://search.kg.ebrains.eu/instances/Project/e39a0407-a98a-480e-9c63-4a2225ddfbe4). Four hemispheres from three post-mortem male Macaca fascicularis monkeys (7.3 ± 0.6 years old; weight 6 ± 0.8 kg) were used in these studies, and coronal brain sections were alternately processed to analyze 14 receptor types: glutamatergic (AMPA, kainate, NMDA), GABAergic (GABA_A_, GABA_B_, GABA_A_ associated benzodiazepine [GABA_A_/BZ]), cholinergic (M_1_, M_2_, M_3_), adrenergic (α_1_, α_2_), serotoninergic (5-HT_1A_, 5-HT_2_) and dopaminergic (D_1_). All experimental protocols followed European Communities Council Directive guidelines for animal care and use. For more information on subjects and procedures, see Impieri et al. (2019) and Niu et al. (2020).

The RC patterns refer to the similarity of the receptor distribution patterns across brain areas. For each brain area, we calculated a representative feature vector consisting of 14 receptor density values. To ensure all receptor types had equal weight for the following analysis, receptor densities were normalized by z-scores within each receptor type. The receptor-based similarity between two areas was measured by computing the Pearson correlation of their representative feature vectors. In this manner, we computed the similarities between each seed area and all target brain areas included in the analysis, resulting in an interareal correlation matrix (N ×M, where N represents the number of seed somatosensory areas and M denotes the total number of cortical areas included; here, N = 34, M = 115).

### Functional connectivity (FC) patterns of macaque somatosensory cortex

The structural and functional MRI data used for FC reconstruction was obtained from PRIME-DE (http://fcon_1000.projects.nitrc.org/indi/indiPRIME.html) (Milham et al., 2018b). Specifically, this dataset anesthetized animals collected at Oxford University, is the largest sample size and the most extensive data (53.33 minutes per animal) (Noonan et al., 2014; Xu et al., 2019) publicly available. The full dataset consisted of 20 rhesus macaque monkeys scanned with no contrast agent on a 3T scanner with a four-channel coil in Oxford. In the present study, one macaque was excluded due to the failure of surface reconstruction, leaving a total of 19 males, aged 4.01 ± 0.98 years, weighing 6.61 ± 2.04 kg. Structural scans were acquired using a T1-weighted MPRAGE sequence (no slice gap, 0.5 × 0.5 × 0.5 mm, TR = 2,500 ms, TE = 4.01 ms, 128 slices), resting-state functional scans were using the following parameters: 36 axial slices, in-plane resolution 262 mm, slice thickness 2 mm, no slice gap, TR = 2,000 ms, TE = 19 ms, 1,600 volumes. For additional details see Noonan et al. (2014).

All data were preprocessed using a Human Connectome Project-like pipeline for Nonhuman Primate as described previously (Autio et al., 2020; Xu et al., 2018; Xu et al., 2019; Xu et al., 2015). For each macaque, the structural preprocessing included denoising, skull-stripping, tissue segmentation, surface reconstruction, and surface registration to align to Yerkes19 macaque surface template (Donahue et al., 2016). The functional preprocessing included temporal compression, motion correction, global mean scaling, nuisance regression (Friston’s 24 motion parameters, white matter, cerebrospinal fluid), band-pass filtering (0.01– 0.1 Hz), and linear and quadratic detrending. The preprocessed data were co-registered to an anatomical space and projected to the mid-thickness cortical surface. Finally, the data were smoothed (FWHM = 3 mm) on the high-resolution native surface, aligned, and down resampled to a 10k surface (10,242 vertices per hemisphere). The preprocessed BOLD activity time courses for each monkey brain were demeaned and then concatenated in time (Fig. 1A).

The FC refers to the statistical correlation of the time series of activation patterns measured in different brain areas. To investigate this, we performed a principal components analysis on activity across all vertices within each area, where the first principal component was taken as the representative activity time course for the area (Rapan et al., 2021; Rapan et al., 2023). Subsequently, we computed the FC between each seed area and all other brain areas by computing the Pearson correlation of their representative activity time course. Following a Fisher’s r-to-z transformation applied to each correlation coefficient, the resulting connection matrix (N ×M, where N represents the number of seed somatosensory areas and M denotes the total number of cortical areas; here, N = 34, M = 115) reflects the FC patterns for all somatosensory-related areas (Fig. 1B).

### Structural connectivity (SC) patterns of macaque somatosensory cortex

The retrograde tract-tracing data used for SC reconstruction stemmed from a subset of an ongoing project by the Team of Henry Kennedy to map the macaque cortical connectome (Froudist-Walsh et al., 2021; Markov et al., 2013; Markov et al., 2014a; Markov et al., 2014b).

Retrograde tracer was injected into a certain cortical area. The tracer was taken up by the axon terminals in this area and retrogradely transported to the cell bodies of neurons that projected to this injected area. The areas where these cell bodies located are source areas.

For a specific injection area, the number of labeled neurons from different source areas varies, ranging from abundant labeling in some areas to minimal labeling in others.

The strength of the inter-areal connection from each source area to the injected area is formulated as a weighted measure using a fraction of labeled neurons (FLN) (Markov et al., 2014a; Markov et al., 2011). This is calculated by dividing the number of labeled neurons in each source area by the total number of labeled neurons across all source areas combined. If a given area had more than one injection, the number of labeled neurons in each source area are summed up across all injections before calculating FLN. For a specific injected area, the number of labeled neurons from different source areas exhibits substantial variability, spanning six orders of magnitude. To account for this, a log10 scale was then applied to FLN with the following formula: log10 (FLN×10^6^ + 1).

Since the tract tracing matrix is partially complete due to the experimental limitation, we applied an imputation model to predict the unknown inter-areal connections based on the known ones. In other words, we inferred connectivity from source area A to injected area B without direct data from B. The imputation method used in this study is adapted from (Molnár et al., 2024), employing a Gradient Boosting Regressor model with the following inputs: the tract-tracing connectivity profile of the source area, the tract-tracing connectivity profile of the target area, and the inter-area distance between the source and target. The profile of a given area A is defined as the comprehensive set of available connectivity information where A serves as the source area.

To improve the model for more precise predictions, we optimized it by incorporating more comprehensive geometrical information and constructing an ensemble of diverse models. The geometrical features added for the optimization include a set of distance profiles, including surface distances on the white matter, mid-thickness, and pial surfaces, as well as a tractography streamline distance profile. The diverse machine learning models used are gradient boosting regressor, multi-layer perceptron and support vector regressor based on polynomial kernel and radial basis function kernel (Vinçon, 2024).

The output of the imputation is the inter-areal connectivity value, which is log10 scaled. To obtain a matrix that best reflects the true connectivity, we hybridized the imputed matrix with the experimental tract-tracing matrix by replacing imputed data with tracer data wherever available. The hybridized matrix is further symmetrized by averaging the connection weights between A and B in both directions.

## Multivariate Statistical Analysis

### Coupling Analyses

To explore the relationship between RC and FC, as well as between RC and SC, we measured RC-FC and RC-SC couplings separately (Fig. 1C). Based on the above analyses, each seed area currently has an RC, an FC, and an SC feature vector. The RC and FC feature vectors consist of 115 elements (i.e., the similarities between this seed area and the 115 target areas in the MEBRAINS atlas), while the SC feature vector consists of 65 elements (i.e., the SC weight between the injected target area and the other source 65 areas in the Lyon atlas (Markov et al., 2014b)).

Since both RC and FC were constructed based on the MEBRAINS atlas, the elements of the RC and FC vectors are the same, the areal RC-FC coupling was measured directly by computing the Pearson correlation between these RC and FC vectors.

The tracer data used to construct the SC were extracted based on the Lyon atlas (Markov et al., 2014b), which differ from the MEBRAINS atlas with respect to terminology, location, and spatial extent of individual brain areas. To enable a comparison between RC and SC, it is necessary to integrate the original tracer and receptor data into a common atlas. For this purpose, we compared delineation in the two atlases across all coronal sections. Our detailed review revealed that although the MEBRAINS atlas showed finer parcels than did the Lyon atlas in frontal and parietal lobes, most of these parcels constitute subdivisions of the areas in the Lyon atlas. Similarly, visual cortex in the Lyon atlas had finer parcels than did the Julich atlas but the former constituted subdivisions of the larger areas in the MEBRAINS atlas. Based on these observations, we adopted the following strategy to create the common atlas: if several small areas (e.g., VIPl, VIPm) in the MEBRAINS atlas can be anatomically merged into a larger area (e.g., VIP) in the Lyon atlas, we integrated these small areas and averaged their receptor densities to represent the receptor densities of the larger area (e.g., VIP) in the common atlas. Similarly, if small areas in the Lyon atlas (e.g., V4fp_LF, V4pc_LF) correspond to subdivisions of a larger area (e.g., V4d) in the MEBRAINS atlas, we averaged the number of labeled neurons in each small area and assigned it as the SC strength of the larger area (e.g., V4d) in the common atlas. In total, 55 areas were transferred into the common atlas following this procedure. Detailed information concerning areal integration is listed in Table S1. Based on this common atlas, we reconstructed the unified SC and RC matrices, with identical elements. Thus, the areal receptor-structure coupling can be measured by computing the Pearson correlation between these unified RC and SC vectors.

### Receptor contribution analysis

We performed the ’leave-one-receptor-out’ method to identify which receptors contribute most to the couplings between RC and FC. The principle of this method is that if a certain receptor promotes the coupling between RC and FC, then coupling decreases when this receptor is removed from the RC vector. Specifically, there were 14 instances of Pearson correlation between RC and FC. At each instance, we removed one receptor type from the original RC vector to obtain a new reduced RC_receptor-_ (e.g., RC_AMPA-_, RC_kainate-_ etc.) vector. We then calculated the Pearson correlation between each reduced RC_receptor-_ vector and the original FC vector to obtain specific receptor-reduced couplings. If the newly measured receptor-reduced coupling is lower than the original one, it means the removed receptor promotes the coupling between RC and FC, and vice versa. Subsequently, the same strategy was performed to identify which receptors contribute most to the RC and SC couplings.

### Hierarchical cluster analyses

To further investigate the roles of RC, FC and SC in the organizational principles of the macaque somatosensory cortex, we identified the grouping patterns among somatosensory areas based on the RC, FC and unified SC matrices, respectively and compared their similarities and differences. Specifically, hierarchical clustering and principal component analyses (PCA) were utilized with Matlab (The MathWorks, Inc., Natick, MA). For the hierarchical cluster analysis, Euclidean distances were computed to represent the similarities between RC/FC/SC of different areas, and the Ward linkage algorithm was chosen as the linkage method. That means, the shorter the Euclidean distance between two areas, the greater the similarity in their RC/FC/SC. The number of stable clusters was determined by a subsequent BIC analysis (Fraley and Raftery, 2007). The PCA allowed multi-dimensional space reduction into two dimensions and visualization inter-areal distances.

## Data availability

The receptor data was accessed from the MEBRAINS Multilevel Macaque Brain Atlas (https://search.kg.ebrains.eu/instances/Project/e39a0407-a98a-480e-9c63-4a2225ddfbe4).

The resting-state MRI data was accessed from the openly available dataset PRIME-DE (http://fcon_1000.projects.nitrc.org/indi/indiPRIME.html). The retrograde track tracing data was accessed from the Kennedy lab (https://core-nets.org).

Any additional information required to reanalyze the data reported in this paper is available from the lead contact upon request.

## Code availability

All code used for data analysis is available at GitHub: https://github.com/MeiqiNiu/somatosensory_stream.git and is publicly available as of the date of publication.

## Acknowledgements

This project has received funding from the European Union’s Horizon 2020 Research and Innovation Programme under the Specific Grant Agreement 945539 (Human Brain Project SGA3), from the European Union’s Horizon Europe Programme under the Specific Grant Agreement No. 101147319 (EBRAINS 2.0 Project), from the Federal Ministry of Education and Research (BMBF) under project number 01GQ1902, from the National Institute of Health (NIH) under grant number R01MH122024-02, from the Deutsche Forschungsgemeinschaft (DFG, German Research Foundation; PA 1815/1-1), from the Helmholtz Association’s Initiative and Networking Fund through the Helmholtz International BigBrain Analytics and Learning Laboratory (HIBALL) under the Helmholtz International Lab grant agreement InterLabs-0015, and the UKRI Biotechnology and Biological Sciences Research Council (BBSRC) via grant number BB/X013243/1. Open Access publication costs are funded by the Deutsche Forschungsgemeinschaft (DFG, German Research Foundation; 491111487). T.X was supported by National Institute of Health (NIH) awards R01MH139349, RF1MH128696, and P50MH109429.

## Author Contributions

**Meiqi Niu:** Investigation, Data Curation, Visualization, Writing – Original Draft, Writing – Review and Editing;

**Seán Froudist-Walsh:** Formal Analysis, Writing – Review and Editing;

**Yujie Hou:** Data Curation, Writing – Review and Editing;

**Ting Xu:** Data Curation, Writing – Review and Editing;

**Lucija Rapan:** Writing – Review and Editing;

**Henry Kennedy:** Data Curation, Writing – Review and Editing;

**Nicola Palomero-Gallagher:** Supervision, Writing – Review and Editing, Project administration.

