## Supplemental information for "Three cortical streams for somatosensory information processing"

Supplementary Table 1. Comparison of brain areas in Jülich Macaque Brain Atlas, Lyon Atlas and a common atlas.

|  |  | Jülich atlas | common atlas | Lyon atlas |  |  |  | Jülich atlas | common atlas | Lyon atlas |
| --- | --- | --- | --- | --- | --- | --- | --- | --- | --- | --- |
| Seed Area | SI | 3al | 3 | 3 |  | Target Area | cing | 31 | 31 | 31 |
|  |  | 3am |  |  |  |  |  | v23 | 23 | 23 |
|  |  | 3ble |  |  |  |  |  | d23 |  |  |
|  |  | 3bli |  |  |  |  |  | p24 | 24 | 24a |
|  |  | 3bm |  |  |  |  |  | a24 |  | 24b |
|  |  | 1 | 1 | 1 |  |  |  | 24 |  | 24c |
|  |  | 2 | 2 | 2 |  |  |  |  |  | 24d |
|  | SII | S2l | S2 | 7op |  |  |  | 32 | 32 | 32 |
|  |  | S2m |  |  |  |  |  | 25 | 25 | 25 |
|  |  | PVl | SII complex | SII |  |  | motor | 4a | M1 | F1 |
|  |  | PVm |  |  |  |  |  | 4m |  |  |
|  |  | PRl |  |  |  |  |  | 4p |  |  |
|  |  | PRm |  |  |  |  |  | F2d | F2 | F2 |
|  | SPL | PEl | SPL | 5 |  |  |  | F2v |  |  |
|  |  | PEla |  |  |  |  |  | F3 | F3 | F3 |
|  |  | PEm |  |  |  |  |  | F4d | F4 | F4 |
|  |  | PEc |  |  |  |  |  | F4s |  |  |
|  |  | PEci | N/A | N/A |  |  |  | F4v |  |  |
|  |  | TSA |  |  |  |  |  | F5d | F5 | F5 |
|  | *ips* | AIP | AIP | AIP |  |  |  | F5s |  |  |
|  |  | VIPl | VIP | VIP |  |  |  | F5v |  |  |
|  |  | VIPm |  |  |  |  |  | F6 | F6 | F6 |
|  |  | PEipe | MIP | MIP |  |  |  | F7d | F7 | F7 |
|  |  | PEipi |  |  |  |  |  | F7i |  |  |
|  |  | MIPd |  |  |  |  |  | F7s |  |  |
|  |  | MIPv |  |  |  |  | PFC | 44 | 44 | 44 |
|  |  | LIPd | LIP | LIP |  |  |  | 45A | 45A | 45A |
|  |  | LIPv |  |  |  |  |  | 45B | 45B | 45B |
|  | IPL | PFop | rostral IPL | 7B |  |  |  | a46d | 46d | 46d |
|  |  | PF |  |  |  |  |  | p46d |  |  |
|  |  | PFG | N/A | N/A |  |  |  | a46v | 46v | 46v |
|  |  | PGop | rostral IPL | 7A |  |  |  | p46v |  |  |
|  |  | PG |  |  |  |  |  | a46df | N/A | N/A |
|  |  | Opt |  |  |  |  |  | a46vf | N/A | N/A |
| Target Area | POJ | LOP | LOP | PIP |  |  |  | p46df | 9_46d | 9_46d |
|  |  | PIP | N/A | N/A |  |  |  | p46vf | 9_46v | 9_46v |
|  |  | DP | DP | DP |  |  |  | 8Ad | 8Ad | 8m |
|  |  | PPt | N/A | N/A |  |  |  | 8Av | 8Av | 8l |
|  |  | PGm | PGm | 7m |  |  |  | 8Bd | 8B | 8B |
|  | visual | V6Adl | V6A | V6A |  |  |  | 8Bm |  |  |
|  |  | V6Adm |  |  |  |  |  | 8Bs |  |  |
|  |  | V6Avl |  |  |  |  |  | 9d | 9 | 9 |
|  |  | V6Avm |  |  |  |  |  | 9l |  |  |
|  |  | V6l | V6 | V6 |  |  |  | 9m |  |  |
|  |  | V6m |  |  |  |  |  | 10d | 10 | 10 |
|  |  | V4dl | V4d | V4fp_LF |  |  |  | 10md |  |  |
|  |  | V4dm |  |  |  |  |  | 10mv |  |  |
|  |  | V4v | V4v | V4fp_UF |  |  |  | 10o |  |  |
|  |  |  |  | V4pc_UF |  |  |  | 11l | 11 | 11 |
|  |  | V3A | V3A | V3A_UF |  |  |  | 11m |  |  |
|  |  | V3d | V3d | V3_LF |  |  |  | 12l | 12 | 12 |
|  |  | V3v | V3v | V3c_UF |  |  |  | 12m |  |  |
|  |  |  |  | V3fp_UF |  |  |  | 12o |  |  |
|  |  |  |  | V3pc_UF |  |  |  | 12r |  |  |
|  |  | V2d | V2d | V2fp_LF |  |  |  | 13a | N/A | N/A |
|  |  |  |  | V2pc_LF |  |  |  | 13b | N/A | N/A |
|  |  | V2v | V2v | V2fp_UF |  |  |  | 13l | 13 | 13 |
|  |  |  |  | V2pc_UF |  |  |  | 13m |  |  |
|  |  | V1d | V1d | V1fp_LF |  |  |  | 14r | 14 | 14 |
|  |  |  |  | V1pc_LF |  |  |  | 14c |  |  |
|  |  | V1v | V1v | V1fp_UF |  |  |  |  |  |  |
|  |  |  |  | V1pc_UF |  |  |  |  |  |  |
|  | auditory | A1 | A1 | CORE |  |  |  |  |  |  |

Supplementary Figure 1

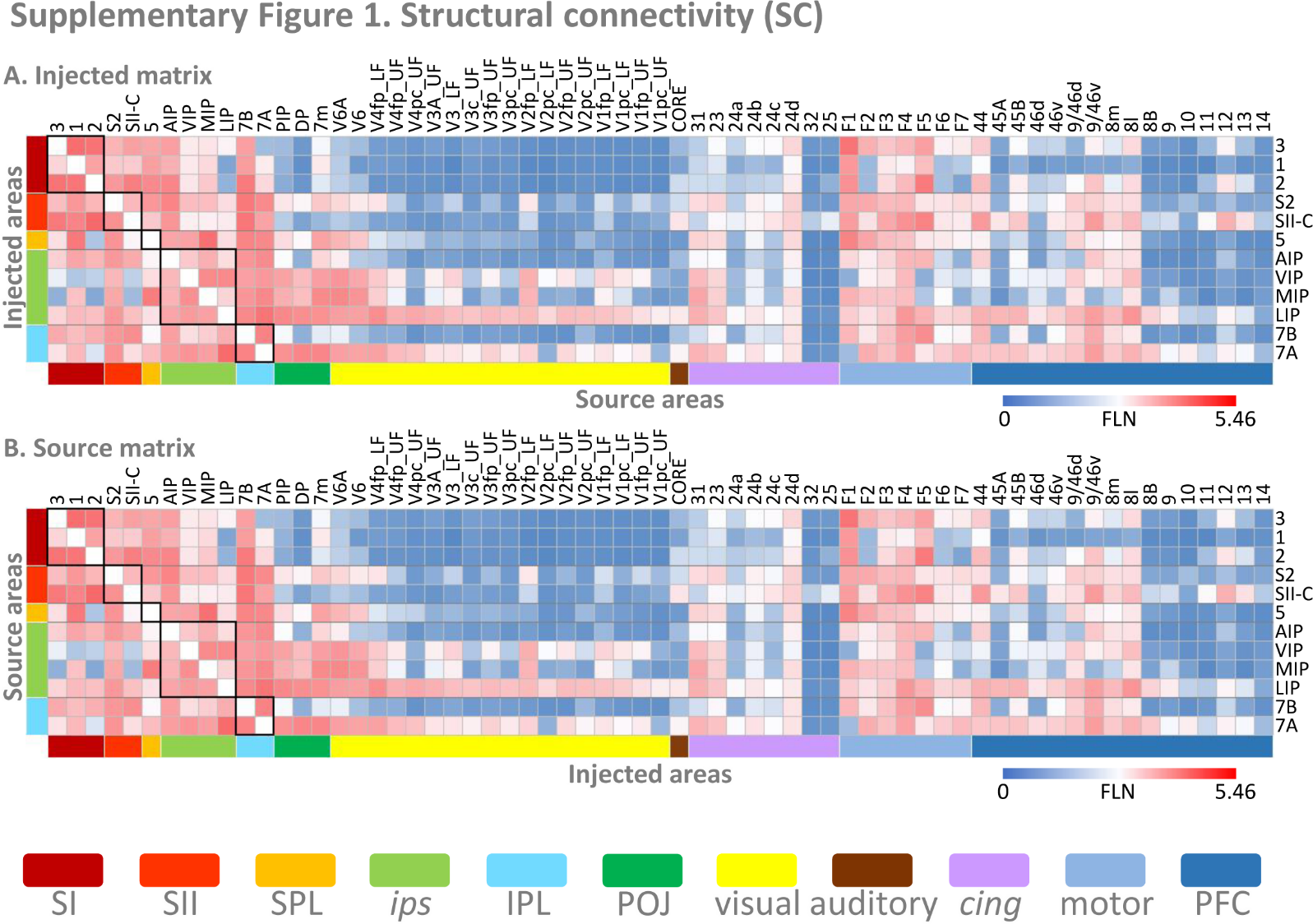

Structural connectivity (SC) matrices of macaque somatosensory cortex. Two asymmetric SC matrices were constructed based on the Lyon atlas. **A**, a 12_target_ × 65_source_ matrix, with somatosensory-related areas as the injected areas. Each row provided the FLN value of input for a given injected area. **B**, a 12_source_ × 65_target_ matrix, with somatosensory-related areas as the source areas. Each row indicates the output FLN value of a source area to other brain areas. The color bars show the FLN strength.
